# Prostaglandin E_2_ production is required for phagocyte CXCR2-mediated skin host defense in obese and hyperglycemic mice

**DOI:** 10.1101/2022.10.02.510554

**Authors:** Nathan Klopsfenstein, Kristin Hibbs, Amondrea Blackman, C. Henrique Serezani

## Abstract

Poorly controlled glucose observed in obese individuals with diabetes is associated with a significantly increased risk of infection, particularly in the skin and soft tissues. *Staphylococcus aureus* is a significant cause of skin and soft tissue infections (SSTIs) in obese and hyperglycemic individuals with growing antibiotic resistance making these infections difficult to treat. However, the events that drive dysregulated skin host defense during hyperglycemia remain to be fully elucidated. Here we examined how the prostaglandin E_2_ (PGE_2_) threshold impacts tissue injury and host defense during methicillin-resistant *S. aureus* (MRSA) skin infection in obese and hyperglycemic mice. Our data show that obesity and hyperglycemia are accompanied by impaired expression of prostaglandin E synthase 1 and PGE_2_ production in infected skin. Restoration of PGE_2_ levels with the PGE analog misoprostol improved infection outcomes in obese and hyperglycemic mice in a manner dependent on E prostanoid 3-mediated cAMP inhibition. Topical misoprostol restored the levels of CXC chemokines and CXCR2+ monocyte and neutrophil recruitment. Here, we are unveiling a defective signaling program that culminates in inadequate CXCR2 phagocyte migration to the infected skin of obese and hyperglycemic mice. Furthermore, these data also lead to a novel drug repurposing opportunity to treat antibiotic-resistant pathogens in hyperglycemic conditions.

## INTRODUCTION

Obesity and hyperglycemia are risk factors for surgical-site infections, nosocomial infections, and skin infections (1–4). Prior case-control studies likewise indicate that overweight and obese individuals are at increased risk of developing cellulitis and skin infections (3). Obesity is also a risk factor for systemic infection, antibiotic treatment failure, and antibiotic resistance due to antibiotic underdosing (3). Furthermore, depending on the antibiotic used, treatment outcomes for certain bacterial pathogens such as *Staphylococcus aureus* may be worse in obese individuals (5). Investigating skin and soft tissue infections in these populations is thus of significant importance due to the rising frequency and severity of pathogens resistant to commonly used antimicrobials, such as methicillin-resistant *S. aureus* (MRSA), which can cause both superficial and systemic infections (5). The increased prevalence of antibiotic-resistant infections, coupled with the rapidly growing incidence of obesity (5), highlights a critical need to determine potential new therapeutic interventions for skin infections in susceptible patient populations.

In patients with hyperglycemia, increased susceptibility to infection has long been correlated to impairments in immune cell effector functions (6–8). Neutrophils and monocytes isolated from the blood of patients with diabetes have shown impaired chemotaxis, phagocytosis, and bacterial killing (7, 9, 10). However, phagocytes from mice and individuals with hyperglycemia also show an increased capacity to produce pro-inflammatory cytokines. However, the mechanisms underlying exaggerated inflammatory response remain to be determined, and the interplay between exaggerated inflammatory response and poor bacterial clearance in people with hyperglycemia is an active area of research in our lab (11–13). While it has been shown that hyperglycemia correlates with impaired immune cell effector functions and worse infection outcomes, the mechanisms driving these defects is an active area of study.

The skin innate immune response to *S. aureus* is orchestrated by skin resident immune and structural cells, including Langerhans Cells (LC), dendritic cells, keratinocytes, and macrophages (14, 15). *S. aureus* recognition leads to the secretion of pro-inflammatory cytokines (IL-1α, IL-1, IFNβ, TNF-α, IL-17, and IL-22), chemokines (CXCL1, CXCL2, and CXCL5), and lipids (leukotriene (LT)B_4_ and prostaglandin (PG) (PGE_2_) to initiate the inflammatory response (15–17).

*S. aureus* skin infection is characterized by initial macrophage-dependent recruitment of neutrophils following the establishment of a localized abscess (15, 18). If the abscess is not tightly organized, bacterial dissemination may occur (15, 19–21) leading to systemic infections. We recently observed that skin infection of hyperglycemic mice results in poor abscess formation, allowing bacteria to grow and disseminate (11, 12). Poor abscess formation in hyperglycemic mice is associated with exaggerated inflammatory response, tissue damage, and uncontrolled bacterial growth (11–13).

We have previously shown an unbalanced production of eicosanoid lipid mediators in the infected skin of lean hyperglycemic mice (11). While we observed an early and sustained increased production of LTB_4_ over time, we detected lower levels of PGE_2_ starting at day 3 post-infection in the skin of hyperglycemic mice. We also demonstrated that decreased local PGE_2_ levels result in poor DC migration to the draining lymph nodes and Th17 generation. When we restored PGE levels with a topical ointment containing the PGE analog misoprostol, we increased dendritic cell migration, Th17 generation and decreased bacterial load in the skin of lean hyperglycemic mice (11).

PGE_2_ is a lipid mediator derived from the metabolism of the cell membrane fatty acid, arachidonic acid, by cyclooxygenase (COX) and the actions of microsomal PGE synthases (MPGES1-2) (22). This prostanoid is abundant at sites of inflammation and influences both pro and anti-inflammatory effects due to actions of different receptors (23–25). The effects of PGE_2_ result from its binding to four distinct cell membrane-associated G protein-coupled E prostanoid (EP) receptors (EP1– EP4) (24, 26, 27). EP1 receptor activates G_q_ (phospholipase C-coupled G protein)-coupled increases in intracellular Ca^2+^; EP3 receptor most often reduces cAMP via inhibitory G protein (G_i_) coupling; and the EP2 and EP4 receptors signal predominantly through stimulatory G protein (G_s_), increasing adenylyl cyclase (AC) activity and subsequent cAMP formation (26, 27). As an inflammatory mediator, PGE_2_ can act as a vasodilator to increase endothelial permeability and enhance cell recruitment (28, 29); however, PGE_2_ actions on phagocytes often inhibit inflammatory cytokine production and limit microbial ingestion and killing by decreasing reactive oxygen species (ROS) production (30–33). The dual role of PGE_2_ and its receptors in modulating the inflammatory response has been observed in several disorders, including diabetes (25).

Given the aberrant inflammatory response observed in the infected skin of hyperglycemic mice, we hypothesized that low PGE_2_ levels may dictate an increased inflammatory response, poor wound healing, and decreased bacterial clearance in obese and hyperglycemic mice. Furthermore, we hypothesized that topical treatment with the FDA-approved PGE analog misoprostol would decrease the amplified inflammatory response and promote wound healing and bacterial clearance in the skin of obese hyperglycemic mice.

Using a diet-induced obesity (DIO) model of hyperglycemia, along with epistatic and gain of function experiments, we unveiled a new role for PGE_2_ and EP3 in driving skin host defense to infections in obese hyperglycemic mice. We found that the impact of misoprostol treatment is dependent on EP3 signaling and CXCR2 actions to improve neutrophil recruitment to the infected skin of obese hyperglycemic mice. Together, these data point to a potential role of PGE_2_/misoprostol as a host-centered therapeutic strategy to treat antibiotic-resistant pathogens during infection in obese and hyperglycemic individuals by directing the inflammatory response to decrease bacterial loads and improve wound healing.

## Methods

### Animals

Mice were maintained according to National Institutes of Health guidelines for the use of experimental animals with the approval of the Vanderbilt University Medical Center (protocol #M1600215) Committees for the Use and Care of Animals. Experiments were performed following the United States Public Health Service Policy on Humane Care and Use of Laboratory Animals and the US Animal Welfare Act. C57BL/6J breeding pairs were initially obtained from the Jackson Laboratory and maintained by breeding at Vanderbilt University Medical Center (VUMC), Nashville, TN, USA. EP3^− /-^ mice (34) were donated by Dr. Richard Breyer (Vanderbilt University Medical Center – VUMC), and CXCR2^fl/fl^_LysM^cre^ (CXCR2^Δmyel^) were donated by Dr. Ann Richmond (Vanderbilt University).

### Streptozotocin (STZ) induced hyperglycemia

For streptozotocin (STZ)-induced hyperglycemia, 6- to 8-week-old male C57BL/6J mice were treated by i.p. injection with 40 mg/kg of STZ (Adipogen) dissolved in 0.1 M sodium citrate buffer once daily for 5 consecutive days (35). Euglycemic control mice received citrate buffer and served as vehicle control. Mice were considered hyperglycemic when blood glucose levels were >250 mg/dL. Mice were rendered hyperglycemic for at least 30 days before MRSA skin infection.

### Diet-Induced Obesity (DIO)

For diet-induced obesity (DIO), 6- to 8-week-old male C57BL/6J mice were placed on a diet containing 60% kcal from saturated fats (Research Diets Inc. #D12492), while lean control mice were fed a nutritionally identical diet containing 10% kcal from saturated fats (Research Diets Inc. #D12450J). Mice were fed for 3 months and examined for body weight and blood glucose levels. Mice were considered hyperglycemic when blood glucose levels were >240 mg/dL.

### *S. aureus* strains and culture

The MRSA USA300 LAC strain was a gift from Bethany Moore (University of Michigan, Ann Arbor, Michigan, USA (36). The bioluminescent USA300 (NRS384 lux) strain was a gift from Roger Plaut (Food and Drug Administration, Silver Spring, Maryland, USA (37). The methicillin-susceptible *S*. aureus (MSSA) Newman strain was donated by Eric Skaar (Vanderbilt University Medical Center, Nashville, TN). MRSA stocks were stored at −80°C and cultured as previously described (38).

### Ointment preparation

Ointments were prepared by emulsifying the active ingredient into 100% petroleum jelly (Vaseline) daily before treatment application. Treatments were applied to cover the infected area with a clean cotton swab. Mice were treated twice a day throughout the infection as indicated.

### *S. aureus* skin infection and treatments

The murine skin infection model was performed as previously described (38). Male mice between 6 and 8 weeks of age were used for *S. aureus* skin infection. Mice were infected with approximately ~3-5 × 10^6^ MRSA CFU subcutaneously, as we have done previously (38). Lesion size was measured daily using a caliper, and the affected area was calculated using the standard equation for the area (length x width) (39). Misoprostol ointment (.03% w/v) was made as described in petroleum jelly (11). To antagonize EP3, mice were treated with an ointment containing the EP3 antagonist L-798, 106 (Tocris Cat. #3342) at .5 mM emulsified in petroleum jelly. Mice were treated topically with an ointment containing either drug or the vehicle control at the indicated time point post-infection and twice daily following initial treatment. For CXCR2 antagonism, mice were injected IP with Navarixin (5mg/kg) (MCE # HY-10198) 3 hours before infection and once daily throughout infection.

### Skin Biopsy Specimens and Bacterial Load

8 mm punch biopsies were collected from naive and infected skin at different time points post-infection and used to determine bacterial counts, histological analysis, cytokine production, mRNA expression, and flow cytometry analysis (38). Skin biopsy samples were collected, weighed, processed, and homogenized for bacterial counts in tryptic soy broth (TSB) media. Serial dilutions were plated on tryptic soy agar. Colony-forming units (CFUs) were counted after incubation overnight at 37 °C and corrected for tissue weight. Results are presented as CFU/g tissue.

### *In situ* mRNA hybridization (RNAscope)

For RNAscope *in situ* hybridization (ISH) analysis, 8 µm skin sections were cut from infection biopsies (40). Images of tissue sections were visualized and acquired using skin sections stained with Hematoxylin and eosin to visualize the infected skin with the Nikon Eclipse Ci and Nikon Ds-Qi2 (Nikon, Tokyo, Japan). To detect *Ptges1 and Ptges2* mRNA expression in FFPE tissues, ISH was performed using the RNAScope Multiplex Fluorescent V2 Assay (Advanced Cell Diagnostics (ACD)) with TSA Plus fluorescein (PerkinElmer) according to the manufacturer’s instructions. Briefly, Advanced Cell Diagnostics designed and synthesized ZZ probe pairs with channel C1 targeting the above mRNA targets (Catalog #497831-PTGES1 and #536661-PTGES2). FFPE tissue sections were co-stained with channel C2 and C3 probes for and *CD68* (ACD Cat. # 316611-C3) mRNA as well. Tissue sections were exposed to ISH target probes and incubated at 40°C in a hybridization oven for 2 hours. After rinsing, the ISH signal was amplified using a company-provided preamplifier and amplifier conjugated to fluorescent dyes (Perkin Elmer #NEL744001KT and #NEL745001KT) as described in the figure legends. Sections were counterstained with 4′,6-diamidino-2-phenylindole (DAPI) (Advanced Cell Diagnostics), mounted and stored at 4°C until image analysis. Image capture was performed using a Keyence BZ-X710 (Keyence, Itasca, U.S.A.), and analysis was performed using HALO Software (Indica Labs). The results are the average of at least 3 different fields of view from at least three different mice/group.

### *In vivo* bioluminescence imaging (BLI) and analysis with IVIS

An IVIS Spectrum/CT (Perkin Elmer) *in vivo* optical instrument was used to image bacterial bioluminescence in the mice. Bioluminescence imaging and analysis were performed as previously described (12).

### Skin Single-Cell Isolation and Staining for Flow Cytometry

Skin biopsies were collected and minced with sterile scissors before digestion in 1 mL of DMEM with 1 mg/mL collagenase D (Roche Diagnostics) for 3 hours at 37℃. Reactions were then quenched with a 10 mM final concentration of EDTA before passing through a 70 μm cell strainer and washed with PBS. Single-cell suspensions were treated with CD16/32 Fc blocking antibodies (Biolegend; catalog 101310; clone 93) to prevent non-specific antibody binding and stained with the fluorescent-labeled antibodies for 20 minutes, followed by fixation using 1% paraformaldehyde. The following antibodies were utilized: F4/80-FITC (Biolegend; catalog 123107; clone BM8), CXCR2-PE (R&D; catalog FAB2164P), Ly6G-PerCP/Cy5.5 (Biolegend; catalog 127616; clone 1A8) Ly6C-AF647 (Biolegend; catalog 128010; clone HK14), CD11b-PE/Cy7 (Biolegend; catalog 101216; clone M1/70). Analyses were completed using FlowJo software (FlowJo, Ashland, OR).

### Detection of Cytokines and Chemokines

Biopsy samples were collected, weighed, and homogenized in TNE cell lysis buffer containing phosphatase and protease inhibitors and centrifuged to remove cellular debris. Skin biopsy homogenates were then analyzed using the pro-inflammatory-focused 18-plex Discovery Assay from Eve Technologies (Eve Technologies, Calgary, AB) to detect cytokines and chemokines. Concentrations for all measurements were corrected for the weight of collected tissue.

### Determination of cAMP levels

Biopsy samples were collected, weighed, and flash-frozen in dry ice. Frozen samples were homogenized in 5-10 volumes (mL solution/gram tissue), based on tissue weight, of 5% TCA. Homogenate was centrifuged at 1500 x g for 10 min, and the supernatant was removed and moved to a clean test tube. TCA precipitates were extracted from the sample by mixing with water-saturated ether and discarding the top layer. To remove residual ether, samples were placed on a heat block at 70℃ for 5 min. Samples were then assayed for cAMP concentration using an ELISA (Cayman Chemical #581001) following manufacturer’s protocol.

### MALDI-MS imaging

Skin biopsies were sectioned at the 12-μm thickness and thaw-mounted onto ITO-coated glass slides. Serial sections were collected for H&E staining. MALDI matrix 9-aminoacridine (9AA) was spray-coated onto the ITO-coated slides via an automatic sprayer (TM Sprayer; HTX Technologies). 9AA was made up of 5 mg/mL in 90% methanol, and four passes were used with a nozzle temperature of 85°C, a flow rate of 0.15 mL/min, 2-mm track spacing, and a stage velocity of 700 mm/min. Nitrogen was used as the nebulization gas and was set to 10-gauge pressure (psig). Images were acquired with a 15T Fourier transform ion cyclotron resonance mass spectrometer (FTICR MS, SolariX; Bruker Daltonics) equipped with an Apollo II dual ion source and Smartbeam II 2 kHz Nd:YAG laser that was frequency tripled to a 355-nm wavelength. Data were collected in the negative ion mode using 2,000 laser shots per pixel with the laser operating at 2 kHz. The pixel spacing was 100 μm (center-to-center distance) in both x and y dimensions. Data were collected from m/z 200–1,400. Tentative metabolite identifications were made by accurate mass, typically better than 1 ppm.

### Detection of eicosanoids by mass spectrometry

Skin biopsy sections were collected and flash frozen with dry ice. Samples were processed by the Vanderbilt University Eicosanoid Core Laboratory as previously described (12). Briefly, skin samples were homogenized and extracted in ice-cold methanol with indomethacin and butylated hydroxytoluene (BHT). Samples were then injected for liquid chromatography-MS (LC-MS). Eicosanoids were identified and quantified based on the mass and amount of known standards.

### Detection of PGE_2_

To measure levels of PGE_2,_ biopsy samples were collected, weighed, and flash frozen. Frozen tissue samples were homogenized in ethanol and centrifuged to remove cellular debris. Samples were then analyzed for the amount of PGE_2_ by EIA (Cayman Chemical # 514010) following manufacturer’s protocol. Concentrations were corrected for the weight of the collected tissue.

### Immunoblotting

Western blots were performed as previously described (41). Protein samples were resolved by SDS-PAGE, transferred to a nitrocellulose membrane, and probed with commercially available primary antibodies against PTGES1 (Invitrogen #PA5-51036) or PTGES2 (Invitrogen #PA5-115806). Membranes were then washed and incubated with appropriate fluorophore-conjugated secondary antibodies (1:10,000, anti-rabbit IgG, IRDye 800CW antibody, #926-32211, Licor). Relative band intensities were quantified using ImageJ software (NIH), as previously described (41).

## Statistical Analysis

Results are shown as a mean +/- SEM and were analyzed using GraphPad Prism 8.0 software (GraphPad Software, San Diego, CA). For comparisons between two experimental groups, a Mann-Whitney test was used, and for comparisons among three or more experimental groups, one-way ANOVA followed by Bonferroni multiple comparison test was used. Two-way ANOVA with repeated measures followed by Bonferroni multiple comparison tests was used to compare infection areas over time between two or more groups. *p*<0.05 was considered significant.

## RESULTS

### Obese and hyperglycemic mice are more susceptible to MRSA skin infection

To investigate the role of diet-induced hyperglycemia on skin host defense, we first tested whether increased susceptibility to MRSA infection is dependent on circulating glucose levels. WT mice were fed a high (HFD) or low-fat diet (LFD) for 12-16 weeks, followed by determination of blood glucose levels. C57BL6/J mice fed an HFD for 3 months showed higher body weight (**S. Fig. 1A**) and blood glucose levels (**S. Fig. 1B**) than mice fed with an LFD. However, we observed that ~40-60% of the HFD mice become hyperglycemic (glucose >240 mg/dL). Therefore, we separated obese mice into euglycemic and hyperglycemic animals to determine whether high glucose differentially influences skin host defense in obese mice. Our data show that while obese and euglycemic mice show increased lesion size and bacterial loads compared to LFD mice during MRSA skin infection, hyperglycemic and obese mice showed significantly larger lesions than both euglycemic obese and LFD mice throughout infection that correlated with increased bacterial loads at day 9 post-infection (**Fig. 1A and B**). Using an IVIS 3D reconstruction of the infection site in both hyperglycemic HFD and euglycemic LFD-fed mice, our data clearly show that infected HFD mice show a deeper and larger infection site when compared to LFD-infected mice (**Fig. 1C**). Next, we performed H&E staining of the infected skin to determine tissue injury and abscess formation. We detected intense recruitment of immune cells to the infected skin of obese hyperglycemic mice, along with large and disorganized abscesses compared to LFD mice (**Fig. 1D**). Given the specific effect of hyperglycemia in enhanced susceptibility to skin infection in obese mice, we decided to focus our studies on comparing obese and hyperglycemic animals with lean animals. Together these data demonstrate that metabolic dysfunction (hyperglycemia) is a key factor in enhanced susceptibility to skin infection in obese mice.

**Figure 1.**
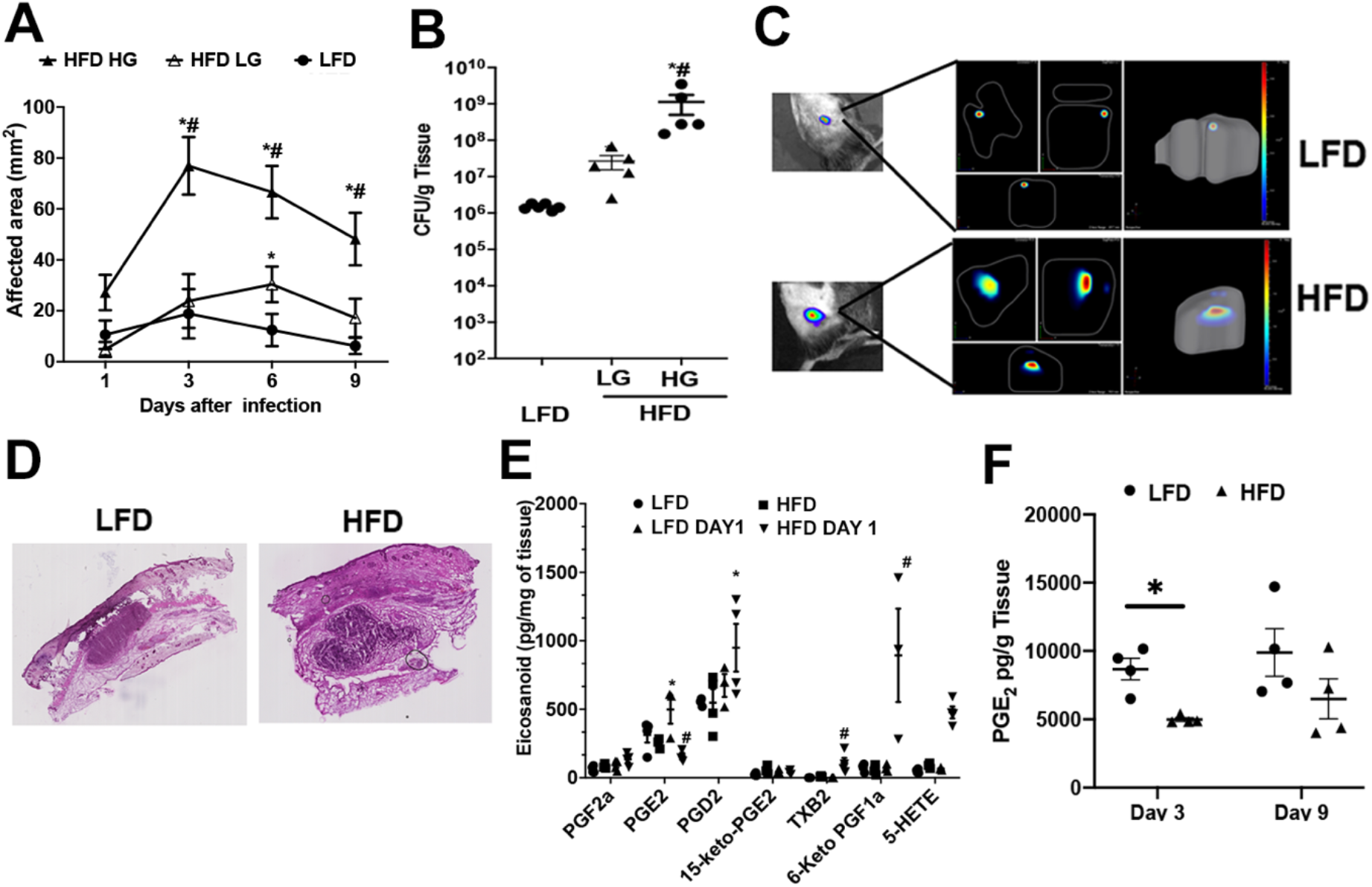
Obese and hyperglycemic mice are susceptible to MRSA skin infection. Male C57BL/6 mice were fed HFD (60 kCal%) and LFD (10 kCal%) for 12 weeks, followed by the determination of blood glucose. **A)** Hyperglycemic and euglycemic obese and lean mice were infected subcutaneously with MRSA (~3 × 10^6^ CFU). Lesion development was monitored every other day for 9 days. Data are mean ± SEM from 5-6 mice from 2–3 independent experiments. *p < 0.05 vs. LFD mice. ^#^p < 0.05 vs. euglycemic obese mice. **B)** Bacterial burden as measured by CFU in skin biopsy homogenates collected from mice as in **A** at day 9 post-infection. Data are mean ± SEM from 5-6 mice from 2–3 independent experiments. *p < 0.05 vs. LFD mice. ^#^p < 0.05 vs. euglycemic obese mice. **C)** (Left) IVIS scanning of bioluminescent MRSA infection in LFD and HFD mice using bioluminescence imaging (BLI) to quantify bacterial burden in the skin on day 9 after infection. (Right) abscess volume of the infected lean and obese mice. Data is representative of at least 5 mice/group. **D)** Representative H&E stained sections of the skin of LFD and HFD mice on day 1 post-MRSA skin infection. Data is representative of at least 5 mice/group. **E)** Eicosanoids were measured by mass spectrometry from skin biopsy homogenates collected on days 1 and 9 post infection from LFD and HFD mice. **F)** PGE_2_ levels in the skin of LFD and HFD mice at day 3 and 9 post infection and measured by EIA. Data are mean ± SEM of at least 3-4 mice from 2 independent experiments. *p < 0.05 vs. LFD mice. ^#^p < 0.05 vs. lean mice.

### Decreased PGE_2_ abundance correlates with increases susceptibility of obese hyperglycemic mice to skin infection

We have previously shown that unbalanced eicosanoid production dictates susceptibility to a skin infection in hyperglycemic mice (11, 12). Here, we investigated the levels of prostanoids and the 5-LO product 5-HETE in the infected skin of obese hyperglycemic and lean mice. We observed that only PGE_2_ levels were reduced at day1 post-infection in infected obese hyperglycemic mice **(Fig. 1E)**. Interestingly, other prostanoids, such PGD_2_, TXB_2_ and PGF1α were increased in the skin of obese mice **(Fig. 1E)**. We further observed decreased PGE_2_ abundance on days 3 and 9 post-infection in HFD compared to LFD controls **(Fig. 1F)**. These data suggest that a specific deficiency in PGE_2_ production could be accounting for inadequate wound healing and skin host defense in obese hyperglycemic mice.

### High glucose impairs PTGES1 expression during MRSA skin infection

Next, we examined the potential cause of lower PGE_2_ production in the infected skin of obese and hyperglycemic mice. Previously, in a model of STZ-induced hyperglycemia, we observed no difference in the expression of enzymes involved in prostanoid production - cyclooxygenase enzymes (COX-1 and COX-2) (42), between euglycemic or hyperglycemic mice. However, whether obesity would influence the expression of COX-1 or 2 during infection remained to be determined. Our data show that skin infection in obese hyperglycemic mice leads to enhanced COX-2 expression **(Fig. 2A)**. However, increased COX-2 expression does not explain lower PGE_2_ levels in the skin of infected obese mice. Therefore, we hypothesized that a failure to induce specific prostaglandin E2 synthases (PTGES1 or PTGES2) contributes to impaired PGE_2_ production in the skin of obese hyperglycemic mice. When we examined gene and protein abundance of PTGES1 and PTGES2 from skin biopsies taken on day 3 post-infection in LFD and HFD-fed mice, we observed a failure to induce PTGES1 in the skin of mice fed the HFD **(Fig. 2A - C)**, with little differences in PTGES2 expression among groups **(Fig. 2A - C)**. These results show that decreased skin PGE_2_ levels correlate with a failure to induce expression of PTGES1 in HFD-fed mice, but not changes in the expression of the constitutively expressed PTGES2. To examine if PGE_2_ synthesis is actively occurring in areas near the abscess and whether macrophages could be involved in deficient PGE_2_ production, we performed *in situ* hybridization against *Ptges1 and Cd68* mRNA. Using biopsy histology samples from LFD and HFD-fed mice at day 3 post-infection, we observed a lower percentage of *Ptges1+* macrophages and total cells in the skin of obese **(Fig. 2D and 2E)** and STZ-induced hyperglycemic mice **(Sup. Fig 2A and 2B)**. Importantly, *Ptges1* mRNA was both located/enriched in the periphery and within the abscess itself **(Fig. 2D)**. These data suggest that hyperglycemia negatively influences PTGES1 expression at both the protein and mRNA level in response to bacterial stimuli in areas of phagocyte recruitment and abscess formation.

**Figure 2.**
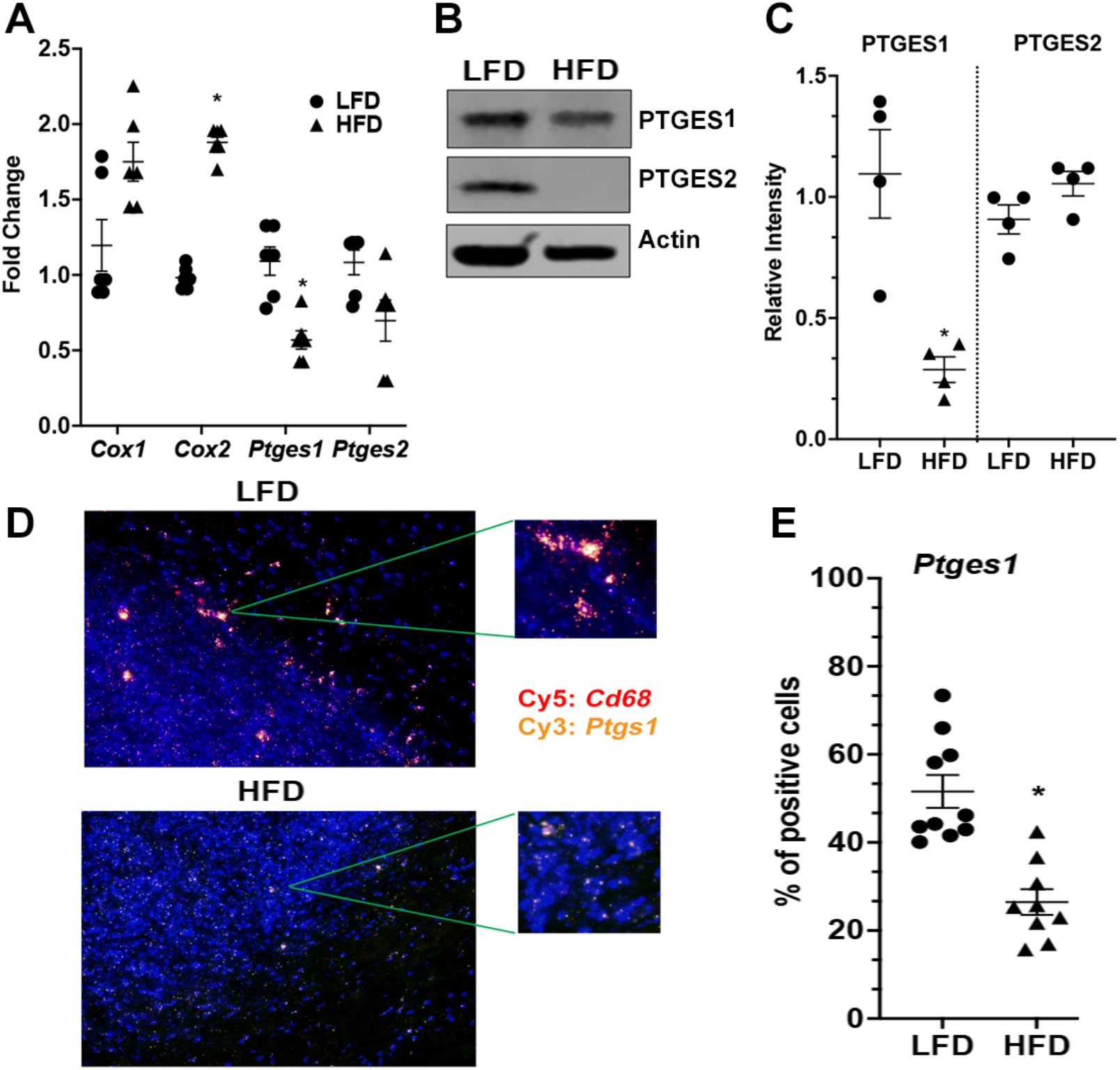
Obesity and hyperglycemia inhibit PTGES1 expression during skin infection. **A)** mRNA expression of the indicated genes from biopsies collected at day 3 post-infection in LFD and HFD fed mice. **B)** Western blotting analysis of PTGES1, PTGES2, and Actin from skin biopsies homogenates collected as in **A. C)** Quantification of densitometry values of the bands shown in **B. D)** Representative RNAscope staining of *Cd68* (red) and *Ptges1* (yellow) in the infected skin of LFD and HFD at day 3 post infection. **E)** Quantification of RNA specks in in at least 10 fields expressed as mean ± SEM of 3-5 mice/group. Each symbol represents the percentage of positive cells in an individual mouse. Data represent the mean ± SEM of 3-10 mice from at least 2-3 independent experiments. *p < 0.05 vs. LFD mice.

### Topical misoprostol improves wound healing and bacterial clearance in different models of metabolic dysfunction

We next sought to examine if increasing local PGE_2_ levels in infected HFD mice would restore skin host defense. We treated HFD and LFD-fed mice topically with the PGE analog misoprostol immediately after MRSA skin infection and twice daily for 9 days post-infection. Initially, we confirmed that topical ointment containing misoprostol is absorbed into the infected skin using IMS **(Fig. 3A)**, which showed misoprostol enriched in areas surrounding as well as inside the abscess. The role of misoprostol in improved skin wound healing and bacterial clearance was evidenced by the fact that misoprostol significantly decreases lesion size and bacterial load when compared to obese hyperglycemic mice treated with vehicle control **(Fig. 3 B - D)**. Furthermore, 3D reconstruction of the bacterial abscess shows that misoprostol also reduces abscess volume within the skin of HFD mice **(Fig. 3E)**. Misoprostol also enhances bacterial clearance and wound healing in methicillin-susceptible *S. aureus* (MSSA) infected obese hyperglycemic mice as well **(S. Fig. 3A)**. Overall, these data point to a critical role for PGE_2_ production in skin host defense that may be contributing to the impaired immune response in obese hyperglycemic mice.

**Figure 3.**
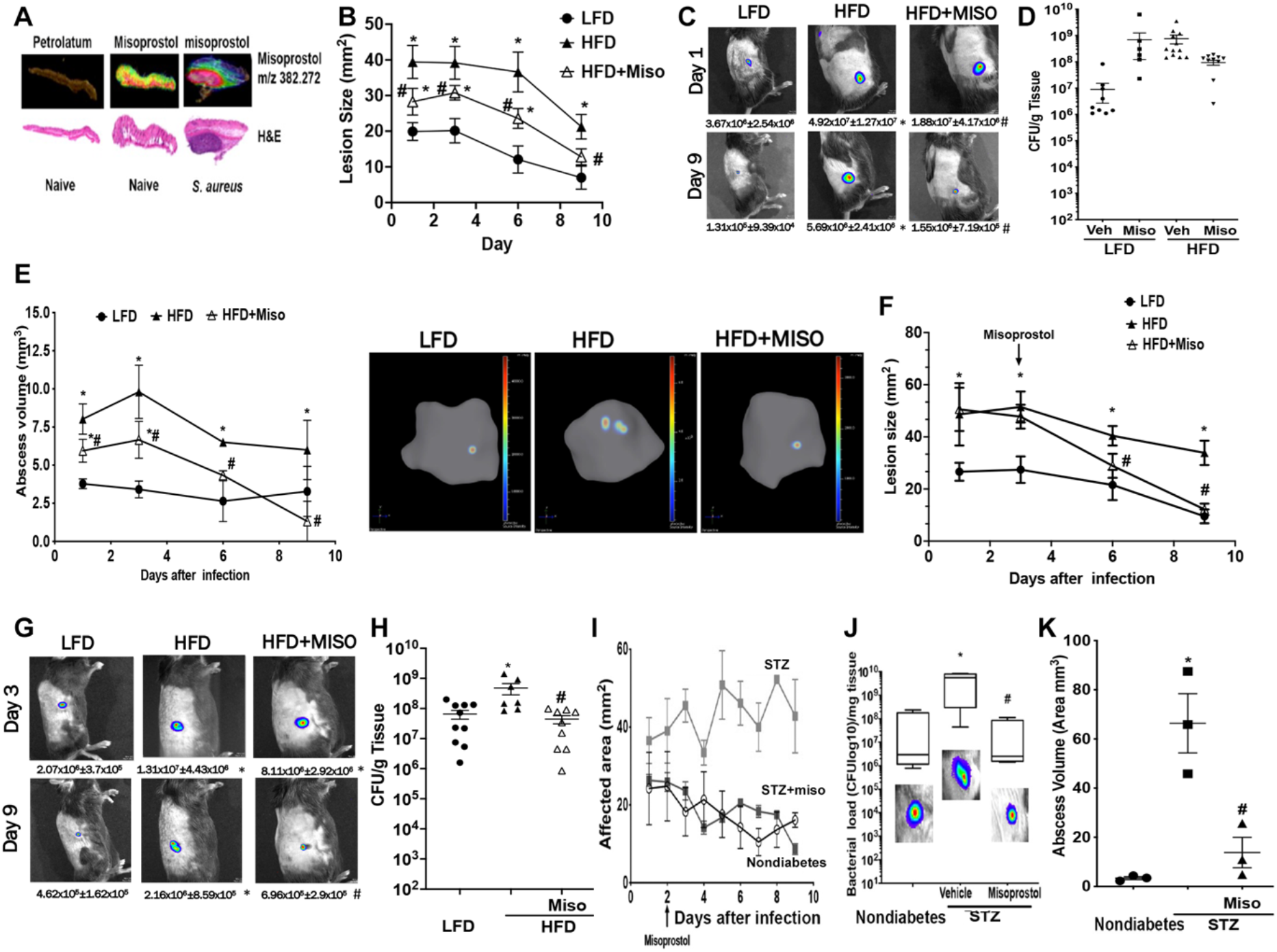
Misoprostol decreases lesion size and bacterial loads in different models of hyperglycemia. **A)** C57BL/6 mice were infected or not with MRSA and treated topically with misoprostol (0.03%) or vehicle control (petrolatum) for 12 h. Top: misoprostol detection using IMS. Bottom: H&E staining from infected and naïve skin. Data are representative of three mice/group. **B)** infection area in LFD and HFD mice infected with bioluminescent MRSA and treated or not with misoprostol as determined using the *in vivo* animal imaging (IVIS Spectrum) detection system. Data are the mean ± SEM of 10 mice from at least two independent experiments. *p < 0.05 vs. LFD mice. **C)** Representative images of bioluminescent MRSA in the skin of mice treated as in **A** using planar bioluminescent imaging with total flux (photons/sec) below. **D)** Bacterial CFUs measured from skin biopsy homogenates from mice in **B** on day 9 post infection. **E)** Left: bioluminescent abscess volume in mice treated as in **B** over time. Right. Representative image of the abscess volume at day 3 post-infection in mice treated as in **B**. Data are representative of at least 5 mice/group. **F)** LFD and HFD mice were infected as in **B**, and 72 h later, HFD mice were treated with or without misoprostol twice daily until day 9 post infection **G)** Representative images of bioluminescent MRSA in the skin of mice treated as in **F** using planar bioluminescent imaging with average flux (photons/sec) below. **H)** Bacterial CFUs measured from skin biopsy homogenates on day 9 post infection of mice treated as in **F. I)** Infection area in control and STZ-induced hyperglycemic mice infected with bioluminescent MRSA and treated as in **F** and measured with a caliper. **J)** Bacterial burden as measured by CFU from skin biopsy homogenates from mice in **I** at day 9 post-infection. Bacterial load was determined by CFU and IVIS (insets). **K)** Bioluminescent abscess volume from mice treated as in **I** on day 3 post infection. Data represent mean ± SEM of 3-12 mice from at least 2 independent experiments. *p < 0.05 vs. LFD or CT mice. #p<0.05 vs. HFD mice treated with vehicle control.

To determine if misoprostol could exert therapeutic effects and improve wound healing and skin host defense in an established infection in obese hyperglycemic mice, we infected LFD and HFD-fed mice and treated with misoprostol topically starting at day 3 post-infection. We observed a significant reduction in the abscess volume, bacterial load, and lesion size in the skin of the misoprostol-treated HFD-fed mice as early as day 6 post-infection **(Fig. 3 F-H)**. Importantly, misoprostol is also effective in eliminating bacteria in established infections in a model of STZ-induced hyperglycemia **(Fig. 3 I-K)**. Together, these data highlight a potential host-derived therapeutic strategy using topical misoprostol to improve wound healing and decrease bacterial loads in patients with chronic hyperglycemia that are susceptible to skin infections.

### Topical misoprostol restores defective chemoattractant production and phagocyte recruitment to the site of infection in obese hyperglycemic mice

Next, we studied how misoprostol impacts the inflammatory milieu in the infected skin of lean versus obese hyperglycemic mice. Using a cytokine bead array, we assessed the abundance of 17 different cytokines and chemokines known to influence host defense and wound healing (15, 16). In the skin of infected obese hyperglycemic mice, we found that IL1β, CXCL1, CXCL2, and CXCL5 levels were decreased compared to infected lean euglycemic mice (**Fig. 4A**). Interestingly, we found that topical misoprostol treatment enhanced the expression of the four chemoattractants, which are known to be critical for neutrophil recruitment in response to skin infection or injury **(Fig. 4A)**. To confirm that the enhanced production of these cytokines/chemokines was not specific to the DIO model of hyperglycemia, we also examined its production in infected, STZ-induced hyperglycemic mice. Examination of infection biopsies from STZ mice revealed parallels between the two models of hyperglycemia with increased IL-1β, CXCL1, CXCL2, but not CXCL5 with misoprostol treatment in infected STZ-induced hyperglycemic mice **(S. Fig. 4A)**.

**Figure 4.**
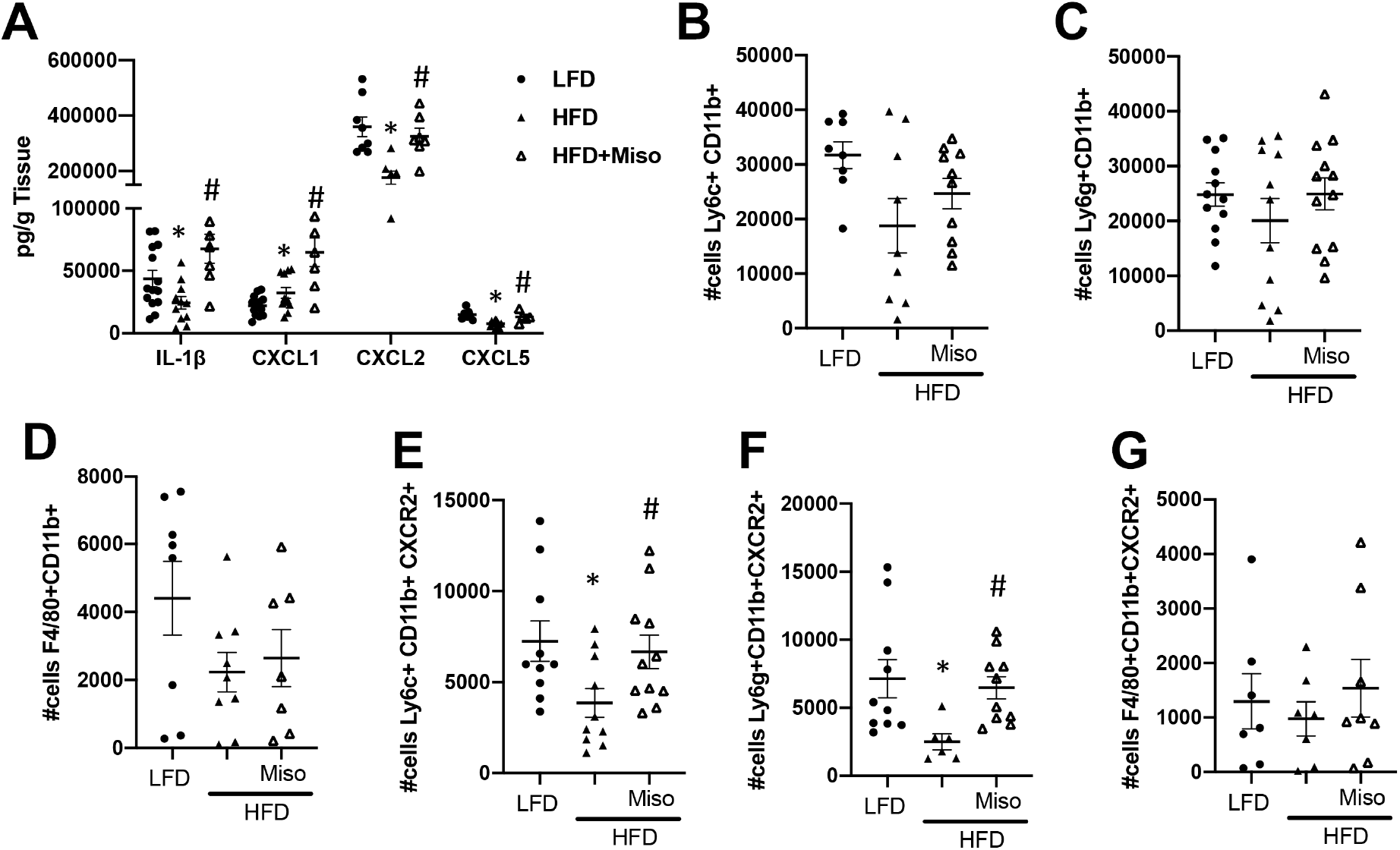
PGE_2_/misoprostol drives CXCR2 phagocyte recruitment to the infected skin of obese mice. **A)** Levels of IL-1, CXCL1, CXCL2, and CXCL5 in skin biopsy homogenates collected at day 3 post infection in LFD and HFD mice treated with or without misoprostol and measured using a bead array multiplex. Absolute cell numbers of **B)** Ly6C+CD11b+, **C)** Ly6G+CD11b+, **D)** F4/80+CD11b+, **E)** Ly6C+CD11b+ CXCR2+, **F)** Ly6G+CD11b+ CXCR2+ and F4/80+CD11b+ CXCR2+ in the infected skin of mice treated as in **A** as detected by FACS. Data are mean ± SEM of 5-10 mice from at least 2 independent experiments. *p < 0.05 vs. LFD mice. #p<0.05 vs. HFD mice treated with vehicle control.

We then aimed to determine whether misoprostol influences phagocyte recruitment to the site of infection in obese hyperglycemic mice. We initially examined the numbers of monocytes (CD11b+Ly6c+), neutrophils (CD11b+Ly6G+), and macrophages (CD11b+F4/80+) in the infected skin of obese and lean mice at day 3 post-infection. We did not detect any significant changes in either monocyte **(Fig. 4B)**, neutrophil **(Fig. 4C)**, or macrophage **(Fig. 4D)** cell numbers among the infection groups. While there was a slight decrease in the overall numbers of monocytes and macrophages in hyperglycemic HFD mice, misoprostol treatment did not appear to have an impact **(Fig. 4B-D)**.

Since topical misoprostol specifically enhances the abundance of chemokines that binds to CXCR1 or CXCR2 (43), we further investigated whether misoprostol influences explicitly the migration of CXCR1/2 expressing phagocytes in the skin of HFD infected mice. We did not observe any effect of misoprostol on the recruitment of CXCR1 expressing monocytes and neutrophils **(Sup. Fig. 5 A and B)**. When we examined the recruitment of CXCR2+ neutrophils and monocytes in the skin of infected hyperglycemic mice treated or not with misoprostol, we observed a significant and specific increase in the numbers of these phagocytes with topical misoprostol treatment compared to vehicle-control treated mice **(Fig. 4E and 4F)**. When we examined these cell populations in the infected and misoprostol-treated skin of STZ-induced hyperglycemic mice, we also observed a specific increase in these same CXCR2+ phagocytes **(S. Fig. 5C and 3D)**. Importantly, we did not see any change in the number of CXCR2+ tissue macrophages in mice treated with misoprostol in either model of hyperglycemia **(Fig. 4G and data not shown)**. These data point to a role for PGE_2_ in promoting targeted cell recruitment to the site of infection during skin infection that may improve not only pathogen clearance but also wound healing and infection resolution.

**Figure 5.**
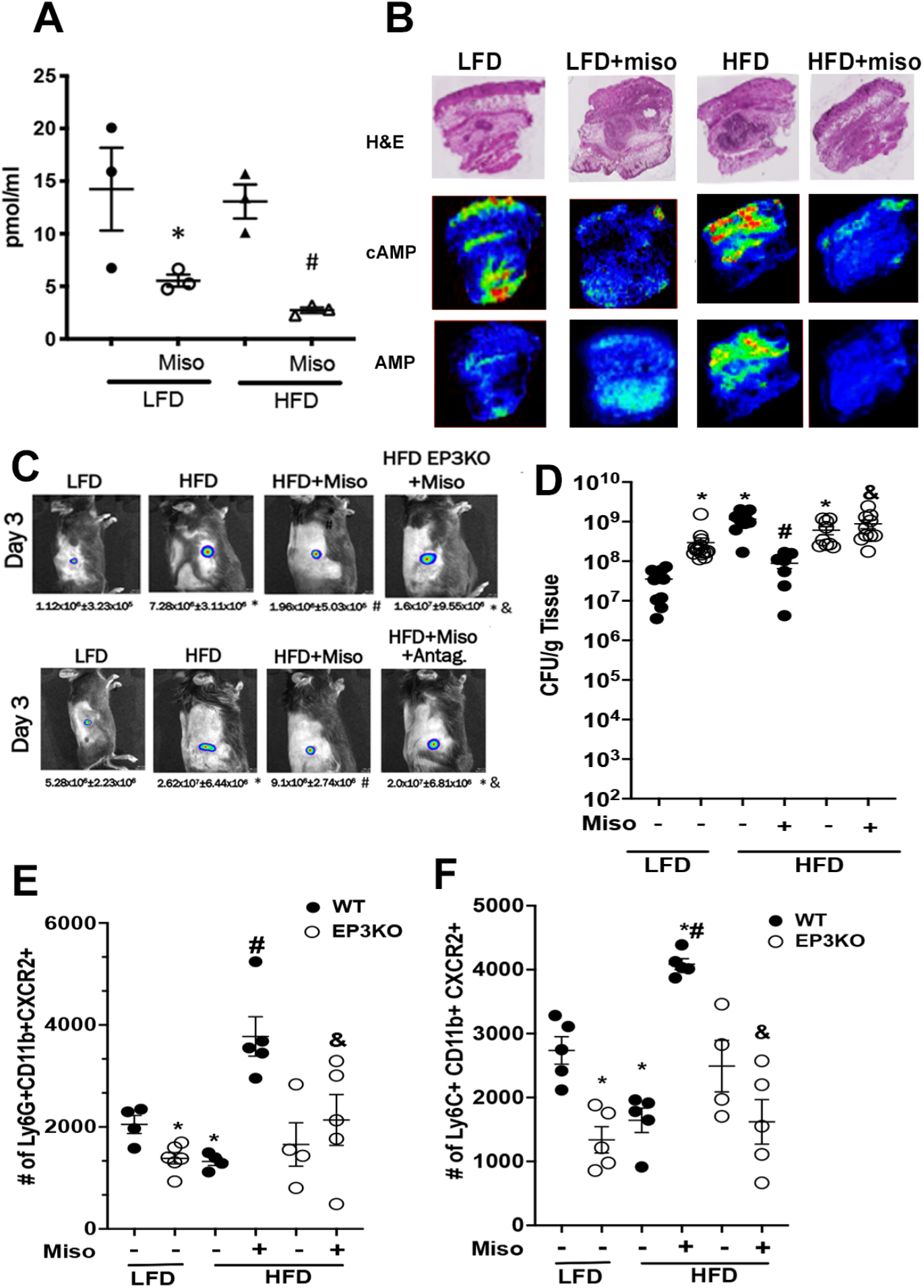
EP3 drives misoprostol-mediated CXCR2 migration to the infected skin of obese mice. **A)** cAMP levels in the infected skin of LFD and HFD mice treated with or without topical misoprostol (0.03%) or vehicle control at day 3 post-infection as measured by EIA. **B)** Top panel: H&E stained sections of the skin of mice treated as in **A** at day 3 post-infection. Bottom panel: MALDI imaging for cAMP and adenosine monophosphate (AMP) determined by imaging mass spectrometry as described in the Methods. Data are representative of at least 5 individual mice/group. **C)** Representative images of bioluminescent MRSA in LFD and HFD mice at day 3 post-infection using planar bioluminescent imaging. **Top**. LFD and HFD mice treated with or without misoprostol plus the EP3 antagonist L-798,106 as described in Methods. **Bottom**: LFD and HFD WT and EP3-/- mice treated with or without misoprostol twice daily for 3 days. Numbers below are quantification of Total flux from mice treated as described above at day 3 post-infection using the IVIS Spectrum **D)** Bacterial burden as measured by CFU from skin biopsy homogenates from LFD and HFD WT and EP3-/- mice treated with or without misoprostol at day 3 post infection **(E and F)**. Absolute cell numbers of **E)** Ly6G+CD11b+ CXCR2+, **E)** Ly6C+CD11b+ CXCR2+ in the infected skin WT and EP3KO LFD and HFD mice treated with or without misoprostol at day 3 post-infection as detected by FACS. Data represent the mean ±SEM from 4-5 mice from 2-3 independent experiments. *p<0.05 vs. LFD. #p<0.05 vs. HFD. &p<0.05 vs. HFD+Miso.

### Misoprostol binds to EP3 to improve wound healing and host defense

We next wanted to determine which EP receptor misoprostol binds to improve wound healing and host defense in obese hyperglycemic mice. As EP receptors are coupled to either G_αs_ or G_αi_, we examined cAMP levels in the skin of infected and misoprostol-treated mice as a readout for EP receptor activation. In skin biopsies from infected LFD and HFD-fed mice treated with or without misoprostol, we observed lower tissue cAMP compared to untreated animals **(Fig. 5A)** as measured via EIA. To assess the cAMP spatial distribution and to confirm that misoprostol treatment lowers tissue cAMP during MRSA skin infection, we also performed IMS on infected biopsies from LFD, and HFD mice treated or not with misoprostol. We observed higher cAMP in the epidermis and the abscess of the HFD-infected mice when compared to cAMP in the infected lean mice **(Fig. 5B)**. Interestingly, misoprostol greatly decreases cAMP in all areas of the infected tissue of obese hyperglycemic mice **(Fig. 5B)**. As EP3 is the sole G_αi_ coupled EP receptor known to lower intracellular cAMP, we hypothesized that EP3 was the main receptor through which misoprostol was driving its therapeutic benefit.

To test this hypothesis, we employed genetic and pharmacological approaches to determine the role of EP3 in skin host defense in obese hyperglycemic mice and in the beneficial effects of misoprostol in skin host defense in obese hyperglycemic mice. LFD and HFD WT and EP3-/- mice were infected and treated with misoprostol immediately after infection twice daily for 3 days **(Fig. 5C and 5D)**. Our data show that EP3 deletion enhances bacterial loads in both obese and lean mice, indicating that EP3 signaling is important to control skin infection under hyper-an euglycemic conditions. Misoprostol did not improve skin wound healing and bacterial load in obese hyperglycemic EP3-/- mice **(Fig. 5C and 5D)**.

To account for possible compensatory effects of whole body EP3 deletion, we also took a pharmacological approach to inhibit EP3 signaling during skin infection and misoprostol treatment. Obese hyperglycemic mice were infected and treated with either misoprostol or misoprostol plus the EP3 antagonist L-798,106 twice daily as we had done previously. After a 3-day infection, we found that misoprostol failed to improve wound healing and bacterial clearance when EP3 was blocked compared to mice treated solely with misoprostol **(Fig. 5C)**. The same experiment in STZ-induced hyperglycemic mice yielded a similar result with EP3 antagonism inhibiting the therapeutic benefit of misoprostol treatment as measured by *in vivo* imaging **(S. Fig 6A and 6B)** and CFU **(S. Fig. 6C)**. Together, these results suggest that EP3 is necessary for the full therapeutic benefit of misoprostol treatment during skin infection in hyperglycemic mice.

**Figure 6.**
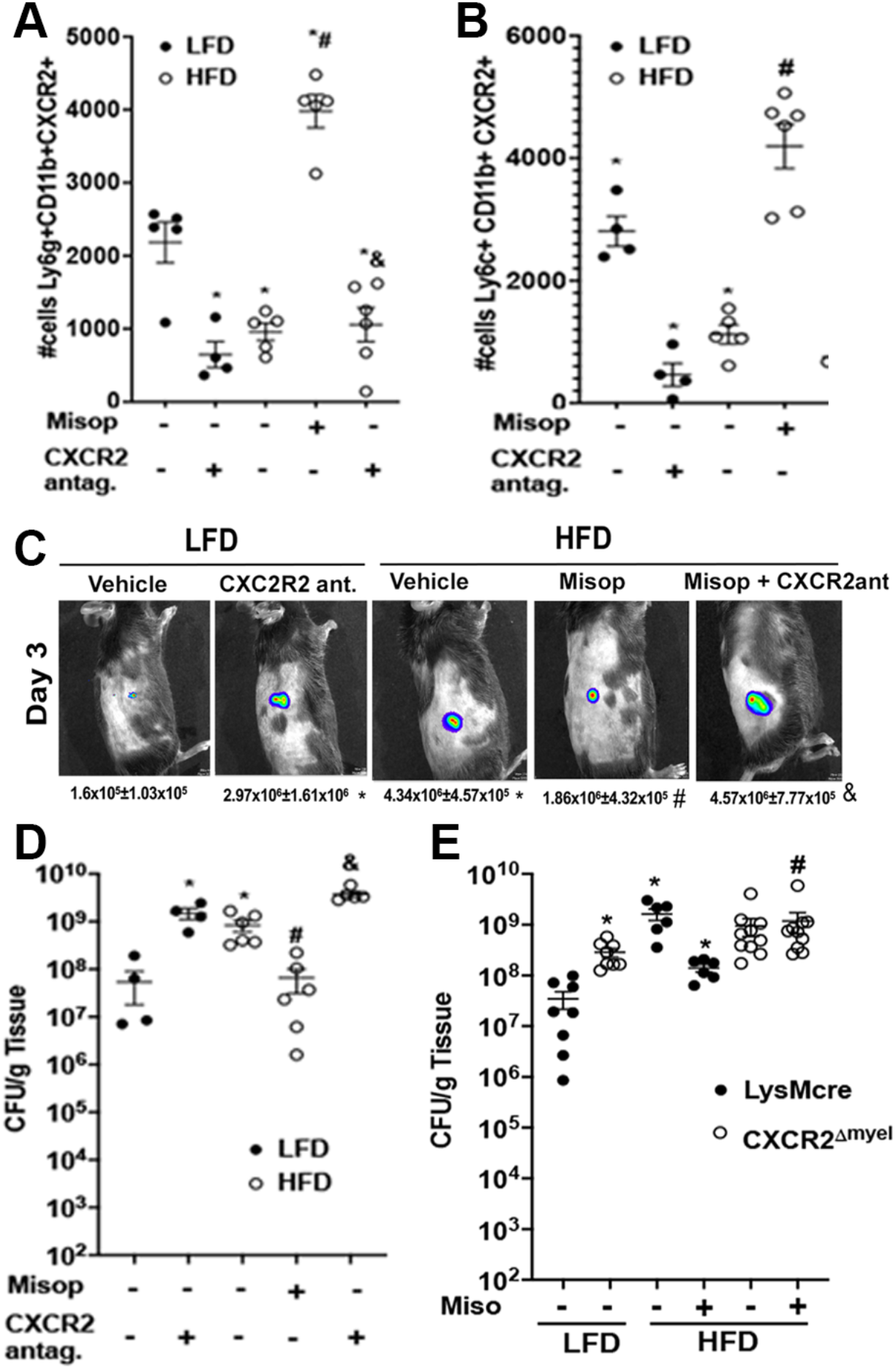
Myeloid CXCR2+ cells are required for topical misoprostol-mediated wound healing and microbial clearance in obese mice. Lean and Obese mice were treated with with topical misoprostol and/or the CXCR2+ antagonist (Navarixin) (5mg/kg) together or individually at day 1 and day 3 post-infection. **A)** Total number of Ly6G+CD11b+CXCR2+ neutrophils in the skin of lean and obese mice treated as in A at day 3 postinfection as measured by flow cytometry. **B)** Total number of Ly6C+CD11b+CXCR2+ monocytes in the skin of lean and obese mice infected and treated as above at day 3 post-infection as measured by flow cytometry. **C)** Representative images of bioluminescent MRSA in the skin of LFD and HFD animals treated with misoprostol or a CXCR2+ antagonist (Navarixin) (5mg/kg) together or individually at day 1 and day 3 post-infection using the IVIS Spectrum. **D)** Bacterial burden as measured by CFU from skin biopsy homogenates from mice treated as above at day 3 post-infection. **E)** Bacterial load in the skin of lean and obese LysMCre and CXCR2^Δmyel^ treated with or without misoprostol twice daily for 3 days determined by CFU Data represent the mean ± SEM from 3-5 mice from 2-3 independent experiments. *p<0.05 vs. LFD. #p<0.05 vs. HFD. &p<0.05 vs. HFD+Miso (One-way ANOVA followed by Bonferroni multiple comparison test).

Next, we investigated if EP3 deletion also impaired the ability of misoprostol treatment to promote CXCR2+ phagocyte recruitment in infected obese hyperglycemic mice. Our data show that misoprostol did not enhance CXCR2+ neutrophils and monocyte recruitment in obese EP3-/- **(Fig. 5E and F)** compared to misoprostol-treated obese WT mice. These data link misoprostol treatment, EP3 activation, and CXCR2+ phagocyte recruitment during hyperglycemic skin infection.

### Misoprostol effects are dependent on EP3-mediated CXCR2 function

Based on the known importance of neutrophils in both pathogen clearance and abscess formation during *S. aureus* skin infection, as well as our data demonstrating improved CXCR2+ immune cell recruitment with misoprostol treatment, we aimed to assess if CXCR2+ cell recruitment was a driving factor in the therapeutic benefit of misoprostol treatment. Therefore, we performed a series of experiments in which we employed genetic and pharmacological approaches to impair CXCR2 actions. we pre-treated LFD or HFD-fed mice with a CXCR2+ antagonist (Navarixin) or vehicle control, followed by misoprostol treatment immediately after MRSA skin infection. Initially, we confirmed that CXCR2 antagonist decreases neutrophil migration in both misoprostol-treated and untreated obese mice both lean and our data show that Navarixin inhibits CXCR2-dependent neutrophil and monocyte migration in both vehicle or misoprostol treated mice (Fig. 6 A and B. Next, we determined the effects of CXCR2 antagonism in bacterial loads in the skin of lean and obese mice. Our data show that tt the end of a 3-day infection, as expected, CXCR2 antagonism increased bacterial loads in lean mice **(Fig. 6C)**, showing a role for CXCR2 in microbial clearance in homeostatic conditions. When we studied the role of CXCR2 in misoprostol effects in obese mice, we found that HFD-fed animals treated with the CXCR2+ antagonist plus misoprostol had significantly worse infection outcomes than those receiving misoprostol alone, as evidenced by CFU **(Fig. 6C)** and IVIS imaging **(Fig. 6D)**. CXCR2 is expressed in migrating myeloid cells and keratinocytes (43). To determine whether misoprostol actions are in keratinocytes or myeloid cells, we treated obese and lean CXCR2^Δmyel^ with misoprostol and showed that: 1) myeloid CXCR2 is indeed essential for skin host defense in lean mice; 2) misoprostol effects are restricted to CXCR2+ myeloid cells **(Fig. 6E)**.

As with our previous results, when this experiment was repeated in an STZ-induced model of hyperglycemic, we observed that mice receiving the combined treatment of misoprostol plus CXCR2 antagonist had significantly worse infection outcomes than untreated animals or those receiving misoprostol alone as measured both by *in vivo* imaging of the infection area **(S. Fig. 7A and 7B)** and CFU **(S. Fig. 7C)**.

To confirm the efficacy of the CXCR2+ antagonist, we performed flow cytometry on infection biopsies collected from these same mice on day 3 post-infection. Upon analysis of these results, we found that animals receiving the CXCR2+ antagonist had significantly reduced numbers of CXCR2+ neutrophils and monocytes **(Fig. 6D and 6E)** compared to untreated animals and prevented the increase seen with misoprostol treatment alone. Similar results were also obtained from flow cytometry analysis of biopsies collected from STZ-induced hyperglycemic mice as well **(S. Fig. 7D and 7E)**, demonstrating our antagonist had the intended effect. Overall, these results indicate a role for CXCR2+ phagocytes in skin host defense against MRSA and identify these cells as a significant contributor to misoprostol-enhanced skin host defense during hyperglycemic skin infection.

## Discussion

People and mice with metabolic dysfunction (obesity and hyperglycemia) are more susceptible to systemic and localized infections caused by different pathogens (3, 5). Enhanced susceptibility to infections correlates with impaired phagocytosis, reactive oxygen species, and microbial killing (3, 44, 45). Also, obesity modifies the T cell responses and antibody formation during infections (44). However, the specific role of hyperglycemia in impaired host defense in obese individuals remains to be determined. We and others have shown that MRSA skin infection in hyperglycemic mice (both STZ-induced and NOD mice models) is accompanied by increased bacterial loads, poor abscess formation, aberrant neutrophil migration, and inflammatory response, leading to tissue injury (11, 12, 45). We have shown that decreased skin PGE_2_ abundance leads to decreased DC maturation and CCR7-mediated DC migration to the lymph nodes and decreased Th17 in the infected skin (11). Here, we move the field forward by filling many gaps by investigating: 1) the cellular players involved in poor bacterial clearance in the skin of obese and hyperglycemic mice; 2) the intracellular events leading to low PGE_2_ production in the infected skin of obese hyperglycemic mice; 3) the therapeutic effects of misoprostol in the skin host defense in obese hyperglycemic mice; 4) the signaling events and cellular targets by which topical misoprostol decreases skin injury and promotes and skin host defense in obese hyperglycemic mice.

Skin host defense is orchestrated by the initial recognition and actions of pathogens by keratinocytes and skin phagocytes (dermal resident macrophages and Langerhans cells) (15, 16). This initial event is crucial to establishing the magnitude and outcomes of the infection. During the homeostatic response, the early events will set the inflammatory response and host defense tone. In conditions where the host is already experiencing systemic alterations in the inflammatory response, the immune cells might respond differently to pathogens, amplifying inflammation and causing damage while failing to control infections. Here, we unveiled that the common ground between STZ and HFD hyperglycemia models, contributing to their impaired skin host defense is the presence of circulating high glucose. The role of high glucose in impaired skin host defense was further confirmed using euglycemic HFD, which showed only slight differences in bacterial loads compared to LFD mice. However, whether dyslipidemia observed in STZ and obese mice could contribute to inadequate skin host defense remains to be determined.

The association between obesity, hyperglycemia and increased risk of skin infection has long been attributed to the immunosuppressive effects of hyperglycemia, an event that has not yet been fully characterized (7, 46–52). In the current proposal, we seek to challenge this dogma. We have generated and published new evidence that obesity and diabetes can also *exaggerate* local immune responses to bacterial infection, resulting in increased lesion size **(Fig. 1A)**, higher bacterial loads **(Fig. 1B)**, increased tissue injury, and failure to form proper abscesses (**Fig. 1D**).

Prior studies have shown that macrophages and neutrophils from obese mice exhibit ineffective bacterial ingestion and reduced production of reactive oxygen and nitrogen species compared to phagocytes from lean mice (4, 49, 53). However, the *in vivo* mechanisms involved in increased susceptibility to skin infections in models of obesity are not well defined. Using genetic models of obesity (ob/ob mice), Farnsworth et al. showed that hyperglycemic and obese ob/ob mice are more susceptible to *S. aureus* hind paw infection, which was correlated with increased neutrophil migration and CXCL1 and CXCL2 production (54). In a different study, Farnsworth et al. also showed that defective antibody production in obese and diabetic mice causes increased skin bacterial load, indicating dysregulated immune responses (45). In 2021, Zhang et al. showed that skin adipocytes from obese mice could not produce antimicrobial peptides and control infection in a TGFβ- and PPARγ activation-dependent manner (44). The fact that adipocytes play an essential role in skin host defense led us to consider that skin lipids might contribute to inflammation and host defense in obese/hyperglycemia and lean/euglycemic mice. However, the role of skin lipids in host defense is not well understood. Our group has started to address this gap in knowledge and has shown a role for the arachidonic acid-derived eicosanoids leukotriene B_4_ (LTB_4_) and PGE_2_ in modulating inflammation and bacterial clearance in hyperglycemic mice (11, 12). Here, we show that skin infection in obese hyperglycemic mice is characterized by robust neutrophil and monocyte migration, poor abscess formation, and increased bacterial loads **(Fig. 1 A-C)**. Interestingly, the phenotype observed in obese mice was also reported in our previous publication of the STZ-induced hyperglycemia model^31^. These data indicate that the inflammatory responses are similarly dysregulated in two independent models of metabolic dysfunction, suggesting that hyperglycemia is a common factor in poor microbial control during skin infection.

In models of sterile wound healing, there is a strong focus on investigating mechanisms of obesity-induced chronic inflammation as the major driving factor of proper healing. Other factors, such as poor revascularization, a major problem in diabetic skin wounds, have also been shown to limit wound healing in DIO mice (48, 55, 56). We observed that increased inflammatory response correlates with lower levels of skin PGE_2_.

The role of PGE_2_ in skin wound healing in mice and people with metabolic dysfunction is somewhat inconclusive. While it has been shown that the levels of PGE_2_ are often elevated in models of sterile wounds in obese mice, infected wounds show decreased PGE_2_ levels in models of hyperglycemia and our model of obesity (57, 58). Importantly, mechanisms underlying increased PGE_2_ levels in sterile wounds point to epigenetic events that increase the abundance of COX2 expression in macrophages. In our model of infection-induced wounds in both obese and hyperglycemic mice, we also observed increased levels of different COX-derived prostanoids, but surprisingly, we detected decreased PGE_2_ abundance in the infected skin of obese hyperglycemic mice. Other investigators have reported lower PGE_2_ production in wound healing models in STZ-induced hyperglycemia in both mice and rats (59, 60). In these models, the authors detected an increased abundance of the prostaglandin transporter PGT (which determines PG catabolism) both *in vivo* and *in vitro* (59, 60). Furthermore, treatment of diabetic mice with the PTG inhibitor T26A increased PGE_2_ levels and restored wound healing. In our model of skin infection in hyperglycemic mice, we did not observe any changes in PGT expression, which could be due to different models of wound healing. Our data show that low PGE_2_ abundance is due to a decreased expression of the terminal enzyme *Ptges1* in infected wounds from hyperglycemic mice as detected by RNAscope analysis in skin sections of obese and lean mice and immunoblotting (**Fig. 2A-**E). Interestingly, we observed decreased PTGS1/PGE_2_ abundance after 3 days post-infection, which suggests that migrated phagocytes, rather than skin resident cells, might be responsible for the decreased PTGS1 expression. Although we did not determine the mechanisms underlying PTGES1 inhibition in the skin of infected obese mice, our data show that culturing bone marrow-derived macrophages from euglycemic mice in high glucose media decreased PTGES1 expression when compared to BMDM cultured in low glucose media, indicating a direct effect of the glucose on the inhibition of PTGS1 expression (Sup. Fig 8). The mechanisms underlying how high glucose influences transcriptional or posttranscriptional events that culminate in PTGS1 inhibition remain to be determined.

Misoprostol, an FDA-approved synthetic prostaglandin E1 analog (PGE1), has mucosal cytoprotective and antisecretory effects and is also used off-label to induce labor. It has been shown that misoprostol inhibits bacterial translocation, modifies bacterial clearance in mice after a burn injury (61, 62), and has cytoprotectant properties against mucosal radiation (61, 62). Misoprostol has multiple uses with different combinations with lidocaine, metronidazole, pentoxifylline, and gentamicin as a wound healing agent for decubitus ulcers or diabetic foot lesions (63).

Misoprostol exerts anti-secretory, anti-inflammatory, and collagen-promoting effects (64–66), by inhibiting thromboxane, MCSF1, and TNF production (64, 67–69). Misoprostol is used on people to heal wounds following radiation therapy, decubitus ulcers, bone and periodontal injuries, and diabetic foot lesions (68, 70, 71). Misoprostol has been found to speed up wound healing in rats but not in dogs in veterinary medicine (71, 72). Here, we described topical misoprostol’s new and potent role in protecting the skin during established MRSA skin infections in obese and hyperglycemic mice.

Misoprostol binds to all EP receptors with different affinities. We have shown that misoprostol binds to EP2 to increase cAMP and inhibit TNF production, microbial ingestion, and killing and production of antimicrobial peptides in rat and murine macrophages (73). The role of EP receptors in skin injury and host defense is not well understood. While most sterile wound healing studies have focused on EP2 and EP4 and cAMP levels, whether EP receptors influence skin host defense remains elusive. Our laboratory has been investigating the mechanisms involved in the high glucose-mediated decrease in cAMP in macrophages. We found increased cAMP production in the infected skin of obese mice compared to infected lean mice (**Fig. 5B**). However, topical misoprostol greatly decreased skin cAMP, suggesting that EP3 signaling is required to improve wound healing and reduce bacterial loads in the infected skin of obese and hyperglycemic mice. The mechanisms underlying EP3-dependent misoprostol effects were not determined, but our *in vitro* data suggest that high glucose inhibits cAMP in MRSA-challenged cells.

When we focused on the role of EP receptors in misoprostol effects, we unveiled a heretofore unknown role for EP3 in both skin host defense and the therapeutic effects of misoprostol treatment. We found that EP3 agonism decreased bacterial loads and increased the production of inflammatory cytokines and chemokines and migration of CXCR2+ phagocytes to the site of infection. These results were not further explored in these manuscripts but are a current research interest of our laboratory. The link between misoprostol, EP3, and CXCR2+ neutrophils in the infected skin of hyperglycemic mice is exciting and unveils a new role for PGE_2_ in the recruitment of neutrophils to the skin. As expected, our data show that myeloid CXCR2 is required for bacterial clearance and neutrophil migration to the site of infection in both lean and obese mice. However, CXCR2 also plays an essential role in the migration of keratinocytes in *in vitro* healing/scratch assays (74). Here, using myeloid-specific CXCR2 deficient mice, we ruled out a potential effect of misoprostol in CXCR2 effects in keratinocyte-mediated wound closure. These findings are interesting since CXCR2 has been implicated in the wound healing of keratinocytes *in vitro* (75). We also assessed whether myeloid CXCR2 and its ligands CXCL1 and CXCL2 influence wound healing during skin infection. Here, we pushed the field forward by uncovering a role for EP3 signaling in promoting the migration of CXCR2+ neutrophils and monocytes. Furthermore, topical misoprostol enhances both CXCL1, CXCL2, and CXCR2+ phagocytes during MRSA skin infection in obese hyperglycemic mice, and such effects are prevented by genetic and pharmacological EP3 blockage. Therefore, misoprostol-mediated EP3 ligation controls different arms involved in CXCR2 activation, including the increased numbers of CXCR2+ cells and its ligands CXCL1 and CXCL2 in the skin of obese mice. The sources of CXCL1/2 during skin infection were not determined.

PGE_2_ released during the resolution phase of the inflammatory response is mainly derived from the efferocytosis of dying cells by phagocytes. These cells also secrete pro-angiogenic factors such as VEGF, which promote revascularization of the tissue during a repair (76). Poor revascularization is another major problem in diabetic and other chronic skin wounds and represents another stage in the wound healing process in which misoprostol treatment may improve skin host defense. Some groups have also demonstrated misoprostol treatment can also promote the release of angiogenic factors from macrophages *in vitro* (77, 78). Since CXCR2+ macrophages and neutrophils, whose recruitment was enhanced with misoprostol treatment, have also been shown to contribute to the process of revascularization, this is an area of interest for future studies in this work. We hypothesize that the duality of both PGE_2_ and misoprostol as initiators and resolvers of the inflammatory response can be beneficial at multiple infection and wound healing stages. This duality also makes misoprostol an ideal therapeutic that can be applied throughout infection to promote different stages of the host immune response and wound healing to improve infection outcomes.

With antimicrobial resistance and the increasing prevalence of obesity/diabetes worldwide, there is a compelling need for safe, inexpensive, non-antibiotic, and host-centered approaches to prevent and treat infections. Here, we provided new fundamental knowledge regarding the regulation of detrimental inflammation to reduce bacterial infection during hyperglycemia using a repurposed FDA-approved drug. We expect that our findings could rapidly translate into clinical use to improve skin infection treatment in obese and hyperglycemic people.

## Acknowledgments

We thank Bethany Moore (University of Michigan, Ann Arbor, MI, USA) for providing the MRSA USA300 LAC strain and Roger Plaut (Food and Drug Administration, Silver Spring, MD, USA) for providing the bioluminescent USA300 (NRS384 lux) MRSA strain. We would like to thank the Serezani laboratory for the input. This work was supported by NIH grants R01HL124159-01 and RAI149207A (to CHS) and T32AI138932 (to NK).

## References

1. Shipman AR, Millington GWM. Obesity and the skin [Internet]. British Journal of Dermatology 2011;165(4):743–750.

2. Sreeramoju P et al. Recurrent skin and soft tissue infections due to methicillin-resistant Staphylococcus aureus requiring operative debridement [Internet]. Am J Surg 2011;201(2):216–220.

3. Huttunen R, Syrjänen J. Obesity and the risk and outcome of infection [Internet]. International Journal of Obesity 2013 37:3 2012;37(3):333–340.

4. Frydrych LM et al. GM-CSF Administration Improves Defects in Innate Immunity and Sepsis Survival in Obese Diabetic Mice [Internet]. The Journal of Immunology 2019;202(3):931–942.

5. Grupper M, Nicolau DP. Obesity and skin and soft tissue infections: How to optimize antimicrobial usage for prevention and treatment? [Internet]. Curr Opin Infect Dis 2017;30(2):180–191.

6. Muller LMAJ et al. Increased Risk of Common Infections in Patients with Type 1 and Type 2 Diabetes Mellitus [Internet]. Clinical Infectious Diseases 2005;41(3):281–288.

7. Alba-Loureiro TC et al. Neutrophil function and metabolism in individuals with diabetes mellitus [Internet]. Brazilian Journal of Medical and Biological Research 2007;40(8):1037–1044.

8. Menne EN et al. Staphylococcus aureus infections in pediatric patients with diabetes mellitus [Internet]. Journal of Infection 2012;65(2):135–141.

9. Foss NT et al. Impaired cytokine production by peripheral blood mononuclear cells in type 1 diabetic patients [Internet]. Diabetes Metab 2007;33(6):439–443.

10. Khanna S et al. Macrophage dysfunction impairs resolution of inflammation in the wounds of diabetic mice [Internet]. PLoS One 2010;5(3):e9539.

11. Dejani NN et al. Topical Prostaglandin e analog restores defective dendritic cell-mediated Th17 host defense against methicillin-resistant staphylococcus aureus in the skin of diabetic mice. Diabetes 2016;65(12):3718–3729.

12. Brandt SL et al. Excessive localized leukotriene B4 levels dictate poor skin host defense in diabetic mice. JCI Insight 2018;3(17). doi:10.1172/jci.insight.120220

13. Filgueiras LR et al. Leukotriene B4-mediated sterile inflammation promotes susceptibility to sepsis in a mouse model of Type 1 diabetes. Sci Signal 2015;8(361). doi:10.1126/scisignal.2005568

14. Bitschar K, Wolz C, Krismer B, Peschel A, Schittek B. Keratinocytes as sensors and central players in the immune defense against Staphylococcus aureus in the skin. [Internet]. J Dermatol Sci 2017;87(3):215–220.

15. Brandt SL, Putnam NE, Cassat JE, Serezani CH. Innate Immunity to Staphylococcus aureus : Evolving Paradigms in Soft Tissue and Invasive Infections. The Journal of Immunology 2018;200(12):3871–3880.

16. Krishna S, Miller LS. Innate and adaptive immune responses against Staphylococcus aureus skin infections [Internet]. Semin Immunopathol 2012;34(2):261–280.

17. Miller LS, Cho JS. Immunity against Staphylococcus aureus cutaneous infections doi:10.1038/nri3010

18. Malachowa N, Kobayashi SD, Lovaglio J, DeLeo FR. Mouse model of staphylococcus aureus skin infection [Internet]. In: Methods in Molecular Biology. Humana Press Inc.; 2019:139–147

19. Kobayashi SD, Malachowa N, Deleo FR. Pathogenesis of Staphylococcus aureus abscesses [Internet]. American Journal of Pathology 2015;185(6):1518–1527.

20. Cheng AG, DeDent AC, Schneewind O, Missiakas D. A play in four acts: Staphylococcus aureus abscess formation [Internet]. Trends Microbiol 2011;19(5):225–232.

21. Singer AJ, Talan DA. Management of skin abscesses in the era of methicillin-resistant Staphylococcus aureus. New England Journal of Medicine 2014;370(11):1039–1047.

22. Park JY, Pillinger MH, Abramson SB. Prostaglandin E2 synthesis and secretion: The role of PGE2 synthases [Internet]. Clinical Immunology 2006;119(3):229–240.

23. Agard M, Asakrah S, Morici LA. PGE2 suppression of innate immunity during mucosal bacterial infection [Internet]. Front Cell Infect Microbiol 2013;4(AUG). doi:10.3389/fcimb.2013.00045

24. Hohjoh H, Inazumi T, Tsuchiya S, Sugimoto Y. Prostanoid receptors and acute inflammation in skin. Biochimie 2014;107(Part A):78–81.

25. Kawahara K, Hohjoh H, Inazumi T, Tsuchiya S, Sugimoto Y. Prostaglandin E2-induced inflammation: Relevance of prostaglandin e receptors. Biochim Biophys Acta Mol Cell Biol Lipids 2015;1851(4):414–421.

26. Sugimoto Y, Narumiya S. Prostaglandin E receptors. Journal of Biological Chemistry 2007;282(16):11613–11617.

27. Biringer RG. A Review of Prostanoid Receptors: Expression, Characterization, Regulation, and Mechanism of Action doi:10.1007/s12079-020-00585-0/Published

28. Martínez-Colón GJ, Moore BB. Prostaglandin E2 as a Regulator of Immunity to Pathogens [Internet]. Pharmacol Ther 2018;185:135–146.

29. Kalinski P. Regulation of Immune Responses by Prostaglandin E 2 [Internet]. The Journal of Immunology 2012;188(1):21–28.

30. Serezani CH et al. Prostaglandin E2 suppresses bacterial killing in alveolar macrophages by inhibiting NADPH oxidase [Internet]. Am J Respir Cell Mol Biol 2007;37(5):562–570.

31. Bourdonnay E, Serezani CH, Aronoff DM, Peters-Golden M. Regulation of alveolar macrophage p40phox: hierarchy of activating kinases and their inhibition by PGE2 [Internet]. J Leukoc Biol 2012;92(1):219–231.

32. Salina ACG, Souza TP, Serezani CH, Medeiros AI. Efferocytosis-induced prostaglandin E2 production impairs alveolar macrophage effector functions during Streptococcus pneumoniae infection [Internet]. Innate Immun 2017;23(3):219–227.

33. Kim SH et al. Distinct protein kinase A anchoring proteins direct prostaglandin E2 modulation of Toll-like receptor signaling in alveolar macrophages [Internet]. J Biol Chem 2011;286(11):8875–8883.

34. Ceddia RP et al. The PGE2 EP3 receptor regulates diet-induced adiposity in male Mice [Internet]. Endocrinology 2016;157(1):220–232.

35. Goyal SN et al. Challenges and issues with streptozotocin-induced diabetes - A clinically relevant animal model to understand the diabetes pathogenesis and evaluate therapeutics. [Internet]. Chem Biol Interact 2016;244:49–63.

36. Domingo-Gonzalez R et al. Prostaglandin E 2 –Induced Changes in Alveolar Macrophage Scavenger Receptor Profiles Differentially Alter Phagocytosis of Pseudomonas aeruginosa and Staphylococcus aureus Post–Bone Marrow Transplant. The Journal of Immunology 2013;190(11):5809–5817.

37. Plaut RD, Mocca CP, Prabhakara R, Merkel TJ, Stibitz S. Stably Luminescent Staphylococcus aureus Clinical Strains for Use in Bioluminescent Imaging. PLoS One 2013;8(3). doi:10.1371/journal.pone.0059232

38. Klopfenstein N et al. Murine Models for Staphylococcal Infection [Internet]. Curr Protoc 2021;1(3). doi:10.1002/cpz1.52

39. Becker REN, Berube BJ, Sampedro GR, Dedent AC, Bubeck Wardenburg J. Tissue-specific patterning of host innate immune responses by staphylococcus aureus α-Toxin. J Innate Immun 2014;6(5):619–631.

40. Klopfenstein N et al. SOCS-1 inhibition of type I interferon restrains Staphylococcus aureus skin host defense [Internet]. PLoS Pathog 2021;17(3):e1009387.

41. Serezani CH, Lewis C, Jancar S, Peters-Golden M. Leukotriene B4 amplifies NF-κB activation in mouse macrophages by reducing SOCS1 inhibition of MyD88 expression. Journal of Clinical Investigation 2011;121(2):671–682.

42. Morita I. Distinct functions of COX-1 and COX-2 [Internet]. Prostaglandins Other Lipid Mediat 2002;68–69:165–175.

43. Zaja-Milatovic S, Richmond A. CXC chemokines and their receptors: A case for a significant biological role in cutaneous wound healing [Internet]. Histol Histopathol 2008;23(11):1399–1407.

44. Zhang LJ et al. Diet-induced obesity promotes infection by impairment ofthe innate antimicrobial defense function of dermal adipocyte progenitors [Internet]. Sci Transl Med 2021;13(577):5280.

45. Farnsworth CW et al. A humoral immune defect distinguishes the response to Staphylococcus aureus infections in mice with obesity and type 2 diabetes from that in mice with type 1 diabetes [Internet]. Infect Immun 2015;83(6):2264–2274.

46. Cluff LE, Reynolds RC, Page DL, Breckenridge JL. Staphylococcal bacteremia and altered host resistance [Internet]. Ann Intern Med 1968;69(5):859–873.

47. Tuazon CU, Perez A, Kishaba T, Sheagren JN. Staphylococcus aureus Among Insulin-Injecting Diabetic Patients: An Increased Carrier Rate [Internet]. JAMA 1975;231(12):1272–1272.

48. Brem H, Tomic-Canic M. Cellular and molecular basis of wound healing in diabetes [Internet]. Journal of Clinical Investigation 2007;117(5):1219–1222.

49. Rendra E et al. Reactive oxygen species (ROS) in macrophage activation and function in diabetes [Internet]. Immunobiology 2019;224(2):242–253.

50. Lipsky BA et al. Skin and soft tissue infections in hospitalised patients with diabetes: Culture isolates and risk factors associated with mortality, length of stay and cost [Internet]. Diabetologia 2010;53(5):914–923.

51. Bowling FL, Jude EB, Boulton AJM. MRSA and diabetic foot wounds: contaminating or infecting organisms? [Internet]. Curr Diab Rep 2009;9(6):440–444.

52. Rio A del, Cervera C, Moreno A, Moreillon P, Miró JM. Patients at risk of complications of Staphylococcus aureus bloodstream infection [Internet]. Clin Infect Dis 2009;48 Suppl 4(SUPPL. 4). doi:10.1086/598187

53. McMurray RW, Bradsher RW, Steele RW, Pilkington NS. Effect of prolonged modified fasting in obese persons on in vitro markers of immunity: lymphocyte function and serum effects on normal neutrophils [Internet]. Am J Med Sci 1990;299(6):379–385.

54. Farnsworth CW et al. Exacerbated Staphylococcus aureus Foot Infections in Obese/Diabetic Mice Are Associated with Impaired Germinal Center Reactions, Ig Class Switching, and Humoral Immunity [Internet]. J Immunol 2018;201(2):560–572.

55. PM S et al. Lack of lymphocytes impairs macrophage polarization and angiogenesis in diabetic wound healing [Internet]. Life Sci 2020;254. doi:10.1016/J.LFS.2020.117813

56. Paulino Do Nascimento A, Monte-Alto-Costa A. Both obesity-prone and obesity-resistant rats present delayed cutaneous wound healing [Internet]. British Journal of Nutrition 2011;106(4):603–611.

57. Liu Z, Benard O, Syeda MM, Schuster VL, Chi Y. Inhibition of Prostaglandin Transporter (PGT) Promotes Perfusion and Vascularization and Accelerates Wound Healing in Non-Diabetic and Diabetic Rats [Internet]. PLoS One 2015;10(7):e0133615.

58. Kämpfer H, Schmidt R, Geisslinger G, Pfeilschifter J, Frank S. Wound inflammation in diabetic ob/ob mice: Functional coupling of prostaglandin biosynthesis to cyclooxygenase-1 activity in diabetes-impaired wound healing [Internet]. Diabetes 2005;54(5):1543–1551.

59. Syeda MM et al. Prostaglandin transporter modulates wound healing in diabetes by regulating prostaglandin-induced angiogenesis [Internet]. Am J Pathol 2012;181(1):334–346.

60. Kampfer H, Schmidt R, Geisslinger G, Pfeilschifter J, Frank S. Wound inflammation in diabetic ob/ob mice: functional coupling of prostaglandin biosynthesis to cyclooxygenase-1 activity in diabetes-impaired wound healing [Internet]. Diabetes 2005;54(5):1543–1551.

61. Topuz Ö, İlhan YS, Doğru O, Aygen E, Sözen S. Effect of melatonin and misoprostol on bacterial translocation in portal hypertensive rats. J Gastroenterol Hepatol 2012;27(3):562–565.

62. Gianotti L, Alexander JW, Pyles T, Fukushima R, Babcock GF. Prostaglandin E1 analogues misoprostol and enisoprost decrease microbial translocation and modulate the immune response. Circ Shock 1993;40(4):243–9.

63. Mahoney J, Ponticello M, Nelson E, Ratz R. Topical misoprostol and wound healing in rats. Wounds 2007;19(12):334–9.

64. Aronoff DM et al. Misoprostol impairs female reproductive tract innate immunity against Clostridium sordellii. J Immunol 2008;180(12):8222–30.

65. Wilson DE. Antisecretory and mucosal protective actions of misoprostol. Potential role in the treatment of peptic ulcer disease. Am J Med 1987;83(1A):2–8.

66. Vandervoort JM, Nieves MA, Fales-Williams A, Evans R, Mason DR. An investigation of misoprostol in the promotion of wound healing. Vet Comp Orthop Traumatol 2006;19(4):191–5.

67. Zaslona Z, Serezani CH, Okunishi K, Aronoff DM, Peters-Golden M. Prostaglandin E2 restrains macrophage maturation via E prostanoid receptor 2/protein kinase A signaling [Internet]. Blood 2012;119(10):2358–2367.

68. Buluç m, Gürdal H, Melli M. Effect of misoprostol and indomethacin on cyclooxygenase induction and eicosanoid production in carrageenan-induced air pouch inflammation in rats. Prostaglandins Other Lipid Mediat 2002;70(1–2):227–39.

69. Mertz-Nielsen A, Eskerod O, Bukhave K, Rask-Madsen J. Misoprostol inhibits gastric mucosal release of endogenous prostaglandin E2 and thromboxane B2 in healthy volunteers. Gut 1995;36(4):511–5.

70. Veness M et al. Use of topical misoprostol to reduce radiation-induced mucositis: Results of a randomized, double-blind, placebo-controlled trial. Australas Radiol 2006;50(5):468–474.

71. Mahoney J, Ponticello M, Nelson E, Ratz R. Topical misoprostol and wound healing in rats. Wounds 2007;19(12):334–9.

72. Vandervoort JM, Nieves MA, Fales-Williams A, Evans R, Mason DR. An investigation of misoprostol in the promotion of wound healing. Vet Comp Orthop Traumatol 2006;19(4):191–5.

73. Zasłona Z, Serezani CH, Okunishi K, Aronoff DM, Peters-Golden M. Prostaglandin E 2 restrains macrophage maturation via E prostanoid receptor 2/protein kinase A signaling [Internet]. Blood 2012;119(10):2358–2367.

74. Sivamani RK. Eicosanoids and Keratinocytes in Wound Healing [Internet]. Adv Wound Care (New Rochelle) 2014;3(7):476–481.

75. Milatovic S, Nanney LB, Yu Y, White JR, Richmond A. Impaired healing of nitrogen mustard wounds in CXCR2 null mice [Internet]. Wound Repair and Regeneration 2003;11(3):213–219.

76. C Y, B H. Cellular Responses to the Efferocytosis of Apoptotic Cells [Internet]. Front Immunol 2021;12. doi:10.3389/FIMMU.2021.631714

77. Gilman KE, Limesand KH. The complex role of prostaglandin E2-EP receptor signaling in wound healing. Am J Physiol Regul Integr Comp Physiol 2021;320(3):R287–R296.

78. Ahluwalia A, Baatar D, Jones MK, Tarnawski AS. Novel mechanisms and signaling pathways of esophageal ulcer healing: the role of prostaglandin EP2 receptors, cAMP, and pCREB. Am J Physiol Gastrointest Liver Physiol 2014;307(6):G602–10.

