## supplemental figs for "Prostaglandin E_2_ production is required for phagocyte CXCR2-mediated skin host defense in obese and hyperglycemic mice"

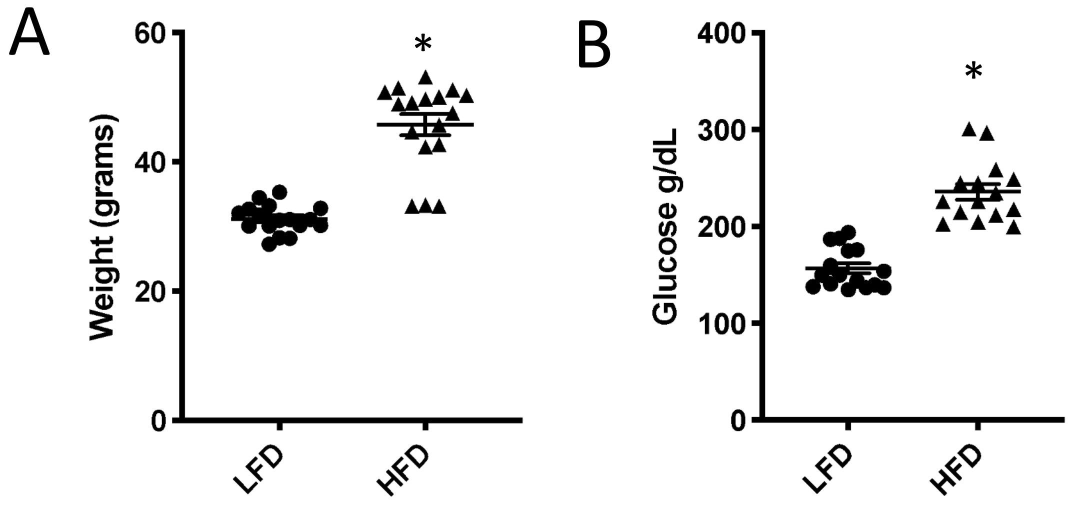


**S. Figure 1. HFD-induced obesity model. A.)** Body weight of C57BL6/J mice placed on a low or high fat diet (LFD or HFD) for 3 months. **B.)** Blood glucose readings from mice in **A.** Data represent the mean ± SEM from 2-3 independent experiments. **p*<0.05 vs. LFD (Mann-Whitney test).

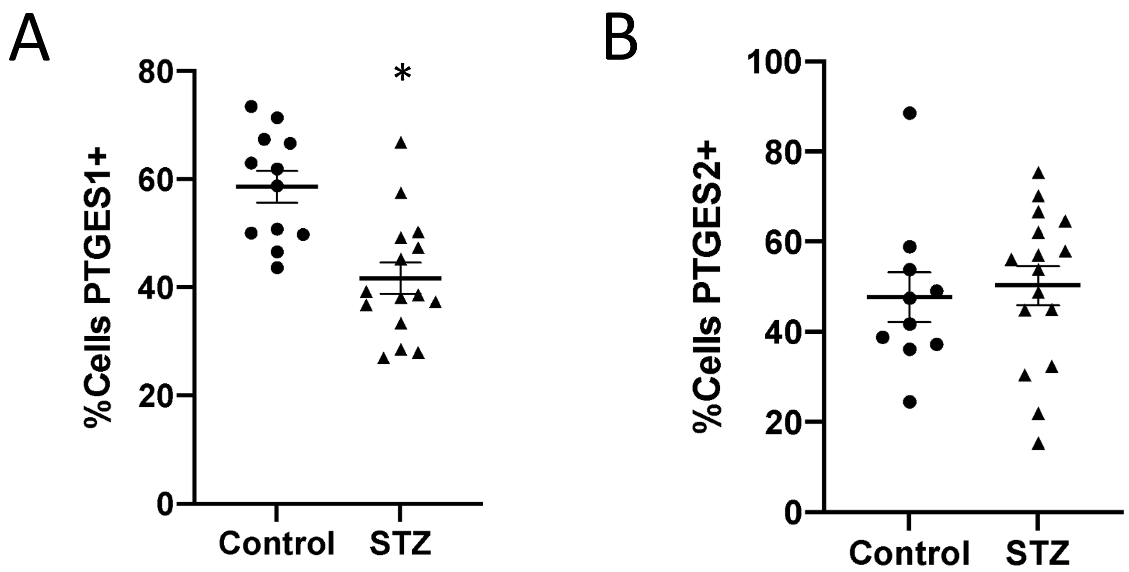


**S. Figure 2. STZ-induced hyperglycemic mice have lower % of *Ptges1*+ cells. A.)** Quantification of the percentage of cells positive for *Ptges1* as determined via *in situ* hybridization quantified utilizing HALO software from biopsies collected at day 3 post-infection in control and STZ-induced hyperglycemic mice. **B.)** Quantification of the percentage of cells positive for *Ptges2* from biopsies as in **A** via HALO software. Data represent the mean ± SEM from 5-12 mice from 2-3 independent experiments. *$p<0.05$ vs. Control (Mann-Whitney Test).

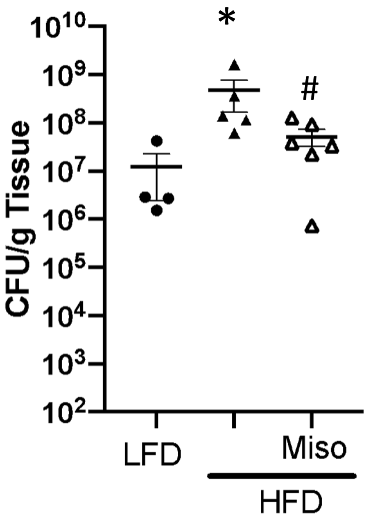


**S. Figure 3. Misoprostol Improves Infection outcome with MSSA Newman in HFD mice.** Bacterial burden as measured via CFU at day 9 post infection in LFD and HFD mice infected with the MSSA Newman strain and treated with or without misoprostol. Data represent the mean ± SEM from 3-6 mice *$p$<0.05 vs. LFD. #$p$<0.05 vs. HFD (One-way ANOVA followed by Bonferroni multiple comparison test).

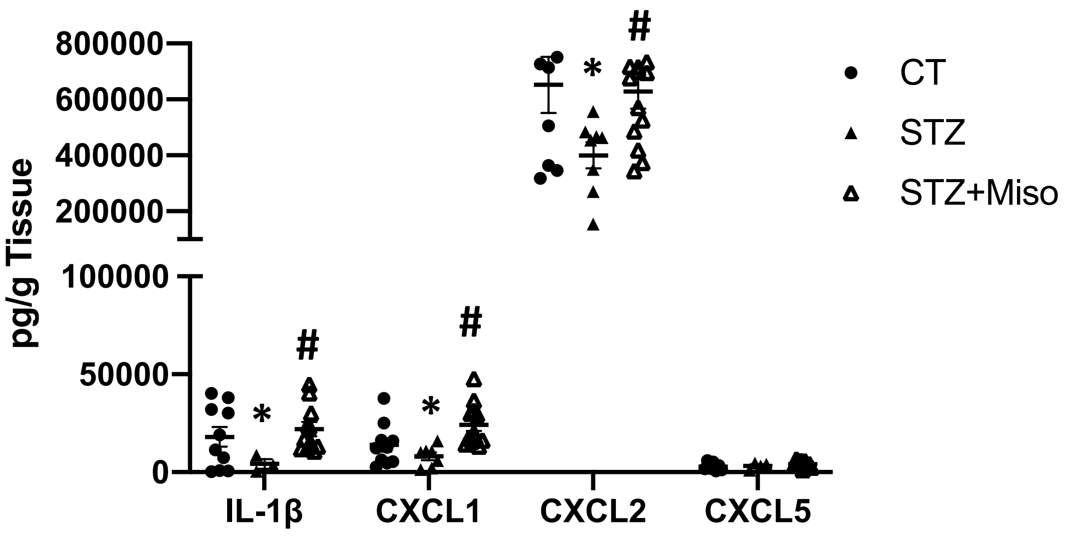


**S.Figure 4. Topical misoprostol increases the production of chemoatractants in STZ-infected mice.** Levels of IL-1β, CXCL1, CXCL2, and CXCL5 in skin biopsy homogenates collected at day 3 post infection in CT and STZ mice treated with or without misoprostol as measured using a bead array multiplex (Eve Technologies). Data represent the mean ± SEM from 5-12 mice from 2-3 independent experiments. *$p<0.05$ vs. CT. #$p<0.05$ vs. STZ (One-way ANOVA followed by Bonferroni multiple comparison test).

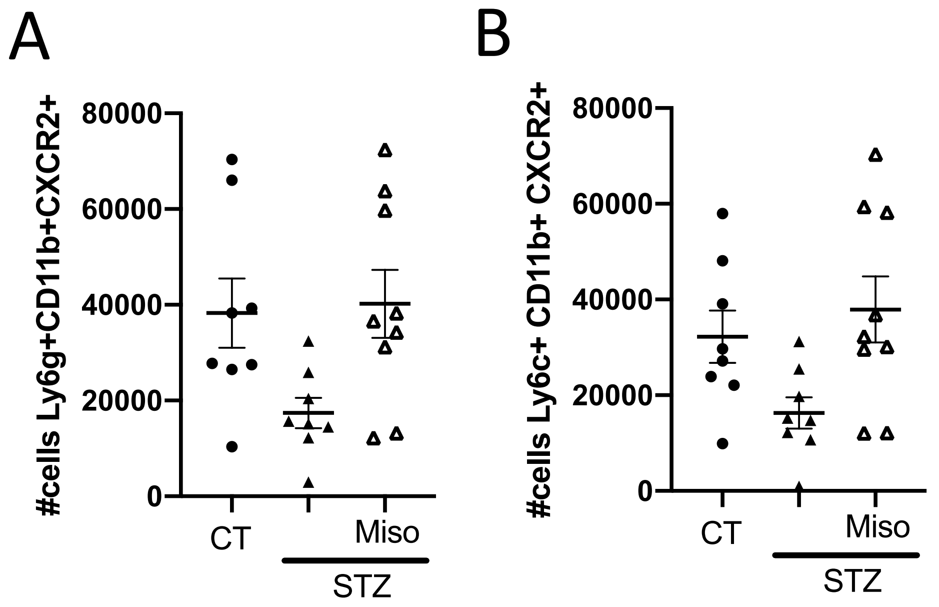


**S. Figure 5. Misoprostol enhances CXCR2+ phagocytes in the skin of diabetic mice**. **A.)** Total number of CXCR2+ neutrophils (Ly6g+CD11b+) in skin biopsies from control and STZ-induced hyperglycemic mice infected s.c. with MRSA and treated with or without misoprostol, at day 3 post-infection. **b.)** Total number of CXCR2+ monocytes (Ly6c+CD11b+) in skin biopsies from mice treated as in **A**. Data represent the mean ± SEM from 8-10 mice from 2-3 independent experiments. *$p<0.05$ vs. CT. #$p<0.05$ vs. STZ (One-way ANOVA followed by Bonferroni multiple comparison test).

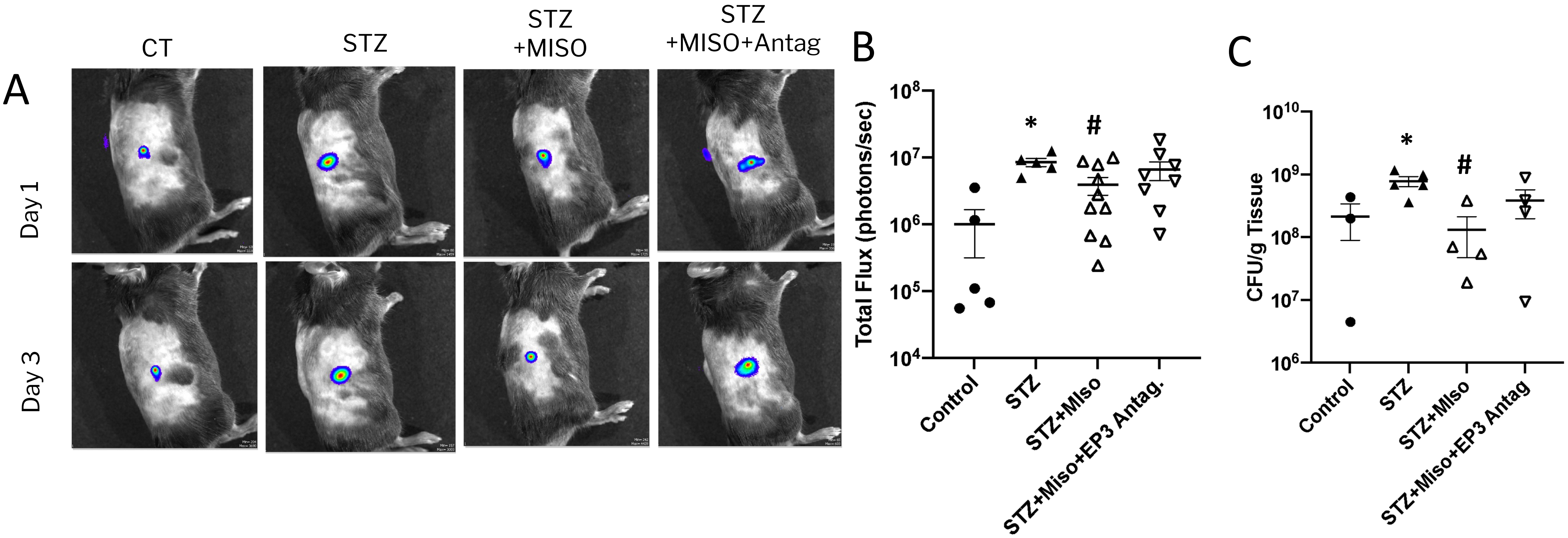


**S. Figure 6. Topical misoprostol improves skin host defense in a manner dependent of EP3 signaling in diabetic mice A.)** Representative images of bioluminescent MRSA in control and STZ-induced hyperglycemic mice treated with or without misoprostol plus an EP3 antagonist (L-798,106) at day 1 and day 3 post-infection using planar bioluminescent imaging. **B.)** Quantification of Total flux from mice treated as in **A** at day 3-post infection using the IVIS Spectrum. **C.)** Bacterial burden as measured by CFU from skin biopsy homogenates from mice treated as in **A** at day 3 post-infection. Data represent the mean ± SEM from 5-6 mice from 2-3 independent experiments. *$p$<0.05 vs. Control. #$p$<0.05 vs. STZ. &$p$<0.05 vs. STZ+Miso (One-way ANOVA followed by Bonferroni multiple comparison test).

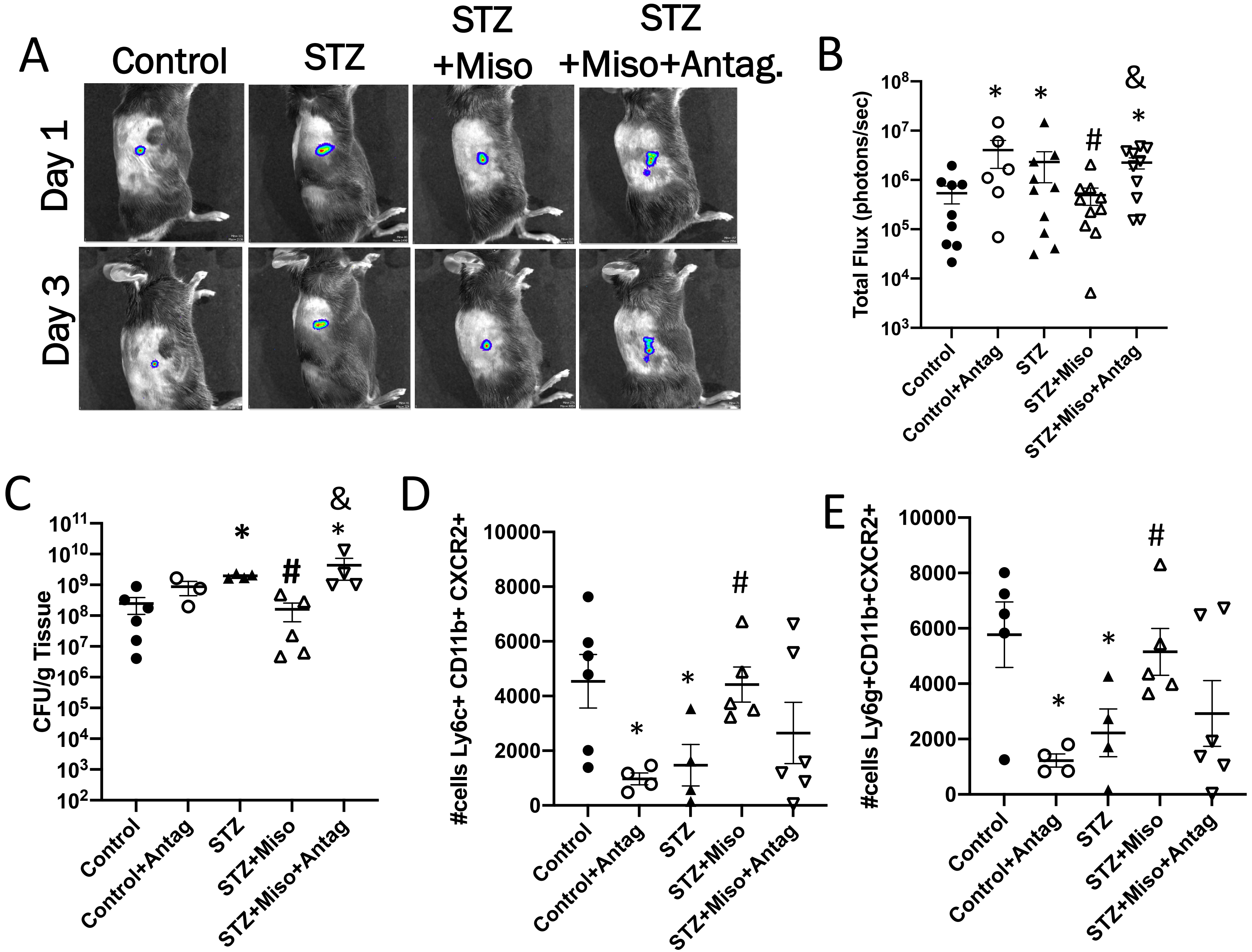


**S. Figure 7.** CXCR2 antagonism blunts therapeutic impact of misoprostol treatment in STZ mice **A.)** Representative images of bioluminescent MRSA in the skin of control and STZ-induced hyperglycemic mice treated with misoprostol or a CXCR2+ antagonist (Navarixin) (5mg/kg) together or individually at day 1 and day 3 post-infection using the IVIS Spectrum. **B.)** Quantification of total flux from bioluminescent MRSA at day 3 post-infection in mice treated as in **A. C.)** Bacterial burden as measured by CFU from skin biopsy homogenates from mice treated as in **A** at day 3 post-infection. Data represent the mean ± SEM from 3-6 mice from 2 independent experiments. *$p$<0.05 vs. Control. #$p$<0.05 vs. STZ. &$p$<0.05 vs. STZ+Miso (One-way ANOVA followed by Bonferroni multiple comparison test).
